# Endocytosis against high turgor pressure is made easier by partial protein coating and a freely rotating base

**DOI:** 10.1101/558890

**Authors:** Rui Ma, Julien Berro

## Abstract

During clathrin-mediated endocytosis, a patch of flat plasma membrane is deformed into a vesicle. In walled cells, such as plants and fungi, the turgor pressure is high and pushes the membrane against the cell wall, thus hindering membrane internalization. In this paper, we study how a patch of membrane is deformed against turgor pressure by force and by curvature-generating proteins. We show that a large amount of force is needed to merely start deforming the membrane and an even larger force is needed to pull a membrane tube. The magnitude of these forces strongly depends on how the base of the membrane is constrained and how the membrane is coated with curvature-generating proteins. In particular, these forces can be reduced by partially but not fully coating the membrane patch with curvature-generating proteins. Our theoretical results show excellent agreement with experimental data.

**SIGNIFICANCE:** Yeast cells have been widely used as a model system to study clathrin-mediated endocytosis. The mechanics of membrane during endocytosis has been extensively studied mostly in low turgor pressure condition, which is relevant for mammalian cells but not for yeast cells. It has been suggested that as a result of high turgor pressure in yeast cells, a large amount of force is needed to drive the progress of the membrane invagination. In this paper, we investigated biologically relevant mechanisms to reduce the force requirement. We highlight the role of boundary conditions at the membrane base, which is a factor that has been largely ignored in previous studies. We also investigate the role of curvature-generating proteins and show that a large protein coat does not necessarily reduce the force barrier for endocytosis.

## INTRODUCTION

Clathrin-mediated endocytosis (CME) is an active process eukaryotic cells use to transport materials from their outside environment to inside of the cell (1–6). During CME, a patch of flat plasma membrane is bent into the cell and severed to release a vesicle (Figure 1a). Deforming the membrane towards the cytoplasm is opposed by membrane’s resistance to bending and membrane tension (7, 8). In walled cells such as plants and fungi, the inward deformation is also opposed by turgor pressure, which pushes the membrane against the cell wall (9–11). In yeast cells, the inner pressure can be up to 1.5 MPa (12, 13). It is conjectured that as a consequence of this high turgor pressure, the membrane invagination exhibits a narrow tubular shape with a diameter of ~ 30 nm in yeast cells (4, 14), while in mammalian cells the invagination exhibits a spherical shape with a diameter of ~ 100 nm due to a relatively low pressure (~ 1kPa) (15).

**Figure 1:**
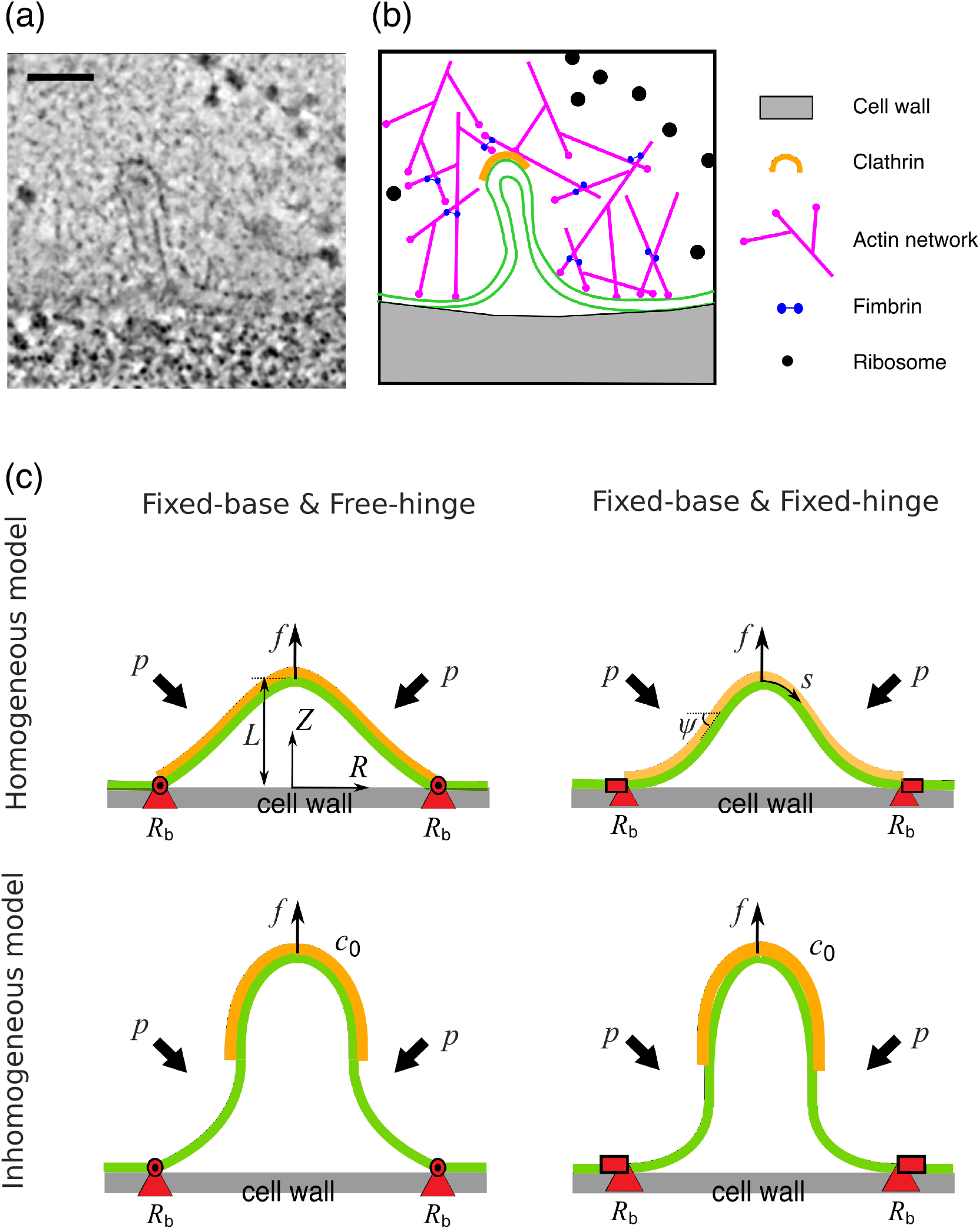
CME in yeast and membrane models for CME. (a) Electron micrograph of a membrane tube formed during CME in budding yeast. The image was obtained from https://www.embl.de/download/briggs/endocytosis.html and adapted under the permission of the authors in (14). The scale bar is 50nm. (b) Schematic illustration of the membrane and key endocytoic proteins shown in (a). The actin network surrounding the membrane tube is depicted as meshwork of branched and crosslinked filaments, though their precise organisation cannot be resolved in the electron micrograph and the meshwork appears as a zone from which ribosomes are excluded. A clathrin coat covering the tip of the membrane tube is also depicted, though the specific spatial distribution of clathrin molecules cannot be resolved in (a). (c) Illustration of the membrane models. The membrane (green layer) is pulled up by a point force *f* against osmotic pressure *p*. The membrane is coated with proteins (orange layer) that locally change the spontaneous curvature of the membrane *c*_0_. The position of the base (red triangles) is maintained at a constant value *R*_b_. We consider a homogeneous model (top) where the membrane is fully coated or fully uncoated with curvature-generating proteins, and an inhomogeneous model (bottom) where the membrane is partially coated. We consider two types of BCs, the free-hinge BC (left) where the membrane is allowed to freely rotate at the base, and the fixed-hinge BC (right) where the membrane angle is fixed.

In the past decade several theoretical models have been proposed to account for the membrane shape evolution during CME (16–21). Most of these models have assumed conditions relevant to mammalian cells, i.e. low turgor pressure (< 1 kPa) and focused on the role of membrane tension. Such tension-dominant membrane deformations have also been extensively studied in *in vitro* experiments where membrane tethers are pulled from giant liposomes (22–24). In contrast, the pressure-dominant regime of membrane deformations, which is relevant to endocytosis in walled cells, has been rarely studied (18). The role of turgor pressure in shaping the membrane has been extensively studied in the case of closed vesicles (25–27). The typical force barrier to invaginate a membrane tube against a membrane tension of 0.01 pN/nm is only 10 – 100 pN, whereas a substantial force (~ 1000 pN) is required to overcome a turgor pressure of 1 MPa (5, 28, 29).

The cytoskeleton protein actin is essential for generating the forces required for CME in yeast cells (10, 30–36). However, the exact organization of actin filaments around the membrane invagination remains elusive. Actin filaments are likely organized into a tight meshwork since ribosomes are excluded from the endocytic sites and actin filaments are heavily crosslinked (Figure1a and b) (37). How the actin machinery produces force to bend the membrane remains unclear. The most commonly accepted hypothesis is that polymerization of actin filaments is converted into a pulling force acting on the tip of the invagination through a push-pull mechanism (28, 38–40). In this mechanism, actin filaments are nucleated on a ring around a patch of clathrin and adaptor proteins. Polymerization of actin filaments at the ring, which is the base of the invagination, pushes the actin meshwork away from the plasma membrane, and in turn pulls the invagination inwards thanks to the adaptor proteins that link actin filaments to the membrane tip.

Membrane can also be bent by proteins that induce membrane curvature. Clathrin molecules can assemble into a cage-like icosahedral lattice composed of hexagons and pentagons *in vitro* (41, 42). The clathrin-lattice alone is able to induce spherical buds from membrane in reconstituted experiments (43). In yeast cells, the clathrin lattice acts as a scaffold linked to the plasma membrane via adaptor proteins and they together form a rigid coat at the membrane invagination tip (44, 45). Based on measurements of the copy number of clathrin molecules in yeast cells, this coat is expected to form a hemi-spherical cap (46). Many clathrin-associated proteins, such as BAR-domain proteins and epsin, have also shown the capacity to induce membrane curvature and might help with CME (47, 48).

In this paper, we study CME under conditions of high turgor pressure and low membrane tension by investigating a theoretical model, which describes how a membrane patch is deformed by a point force and by proteins that induce membrane curvature. In the absence of coat proteins, we show that as a result of high turgor pressure (1 MPa), a large amount of force is needed to merely start deforming the membrane and an even larger force is needed to pull a membrane tube. We also show that the magnitude of these forces strongly depend on the constraints at the base of the membrane patch. In particular, the force to start deforming the membrane increases with the base radius, while the force barrier to pull a membrane tube decreases with the base radius. The forces also depend on whether the angle of the membrane at the base can freely rotate or not.

When the membrane is coated with curvature-generating proteins, we show that the forces to deform partially coated membranes are quantitatively and qualitatively different from the forces to deform fully coated membranes. By partially coating the membrane, the force barrier that is usually present for fully coated membranes can be dramatically reduced to zero, which implies that the membrane can be spontaneously curved up into a vesicular shape.

We find excellent agreement between our theory and experiments. With a single set of parameters for the partially coated membrane model, we can fit geometric features of the membrane shape obtained via electron tomography across different stages of CME. From the comparison, we estimate that the force required for CME in yeast cells is ~ 2500 pN if the membrane angle at the base is free to rotate. This result suggests that actin polymerization alone is insufficient to provide the force to drive the membrane invagination during CME.

## METHODS

### Model of the membrane patch at the endocytic site

We model the membrane patch at the endocytic site as an axisymmetric two-dimensional surface. The shape of the membrane is parameterized with its meridional coordinates [*R*(*s*), *Z*(*s*)], where *s* is the arclength along the meridional direction (Figure1c). The angle *ψ*(*s*) spans between the tangential direction and the horizontal direction. The actin polymerization force is modeled as a point force *f* acting at the symmetric center of the membrane, which is lifted to a height *L* relative to the cell wall (Figure1c). The membrane patch is in contact with the cell wall at a base radius of *R*_b_, which is covered by a ring of proteins as observed in recent experiments (28). We assume the proteins serve as anchors that fix the base of the membrane to the cell wall, therefore *R*_b_ is a constant. Outside of *R*_b_ there is a lipid reservoir such that the membrane tension *σ* is kept constant at the base points. An isotropic turgor pressure *p* is exerted on the membrane, which possesses a bending rigidity *κ* and spontaneous curvature *c*_0_ due to protein coating. Here we assume the turgor pressure *p* is a constant and neglect the concentration change caused by volume reduction upon endocytosis because the reduced volume of the membrane invagination only occupies a tiny fraction (1 /10^6^) of the total cell volume. The free energy of the membrane, which takes into account of the influence of curvature-generating proteins, can be written as

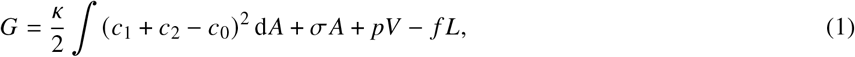

where *c*_1_ and *c*_2_ denote the two principle curvatures of the membrane surface (49), *A* denotes the surface area and *V* denotes the volume between the membrane and the cell wall. The reference state for the free energy *G* in Eq. (1) is a vertically flat and horizontally circular shape. We consider both a homogeneous model where the spontaneous curvature *c*_0_ is spatially uniform - such as a bare membrane or a membrane fully coated with curvature-generating proteins - and an inhomogeneous model where *c*_0_ is spatially varied - such as a membrane partially coated by curvature-generating proteins (Figure1c).

Due to rotational symmetry about the *z*-axis, the free energy of the membrane in Eq. (1) can be expressed as a functional

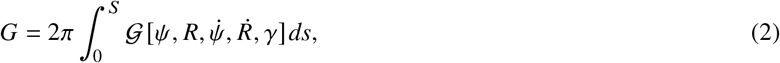

where 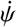 and 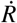 denote their derivatives with respect to the arclength *s*, *S* denotes the total arclength from the tip to the base, *γ* is a Lagrangian multiplier that enforces the geometric relation 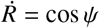 (see Appendix for the explicit form of 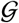). The shape of the membrane is determined by minimization of the free energy *G* with respect to small variations of the membrane shape variables *δψ* and *δR*. Proper boundary conditions (BCs) at the base, where the ring of proteins are formed and the membrane is in contact with the cell wall, are also needed to determine the membrane shape. The exact BCs require knowledge of the microscopic interactions between the membrane, the cell wall, and the ring of proteins. As these microscopic interactions are unclear, we choose to derive the BCs in the following way. The small variations of *δψ* and *δR* result in variation of the free energy *δG*, which consists of boundary terms like 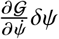 and 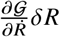. Four types of BCs at the base can be identified by letting these boundary terms vanish (Table 1). Physically they correspond to the combination of whether the base radius is fixed or variable, and whether the angle of the membrane at the base is fixed or free to rotate. We focus on the two BCs where the base radius is fixed (*R = R*_b_) and refer them as free-hinge BC (BC1 in Table 1) if the membrane angle is free to rotate 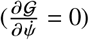 and fixed-hinge BC (BC2 in Table 1) if the membrane angle is fixed to zero (*ψ* = 0). We also compare our results with a previous work (18), which studied the homogeneous model with a BC where the base is free to move and the membrane angle is fixed (BC4 in Table 1).

**Table 1:** Possible boundary conditions at the base of the endocytic membrane.

|  | Base radius | Membrane angle at the base | Mathematical definition |
| --- | --- | --- | --- |
| BC1<sup>a</sup> | fixed | free | $R = R_b, \frac{\partial \mathcal{G}}{\partial \dot{\psi}} = 0$ |
| BC2<sup>b</sup> | fixed | fixed | $R = R_b, \psi = 0$ |
| BC3 | free | free | $\frac{\partial \mathcal{G}}{\partial \dot{R}} = 0, \frac{\partial \mathcal{G}}{\partial \dot{\psi}} = 0$ |
| BC4<sup>c</sup> | free | fixed | $\frac{\partial \mathcal{G}}{\partial \dot{R}} = 0, \psi = 0$ |

<sup>a</sup> Referred to as the free-hinge BC.
<sup>b</sup> Referred to as the fixed-hinge BC.
<sup>c</sup> BC4 has been studied in (18) for the homogeneous model.

## RESULTS

### The characteristic forces to elongate a membrane tube are different between pressure-dominant and tension-dominant conditions

To demonstrate the distinct physics of CME between pressure-dominant and tension-dominant conditions, we approximate the elongated endocytic invagination (as in Figure 1a, for example) as a cylindrical tube of radius *R* and length *L* and derive analytic formulas for the forces to elongate a membrane tube. The free energy (1) under this approximation becomes

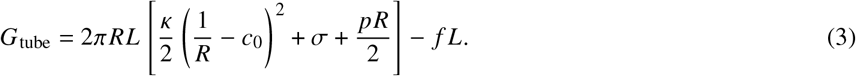

Without considering the effect of the spontaneous curvagture (*c*_0_ = 0), in the case of pressure-dominant condition (*σ* = 0), by minimization of *F*_tube_ with respect to *R* and *L*, we obtain the characteristic tube radius *R*_p_ and the corresponding force *f*_p_ (18):

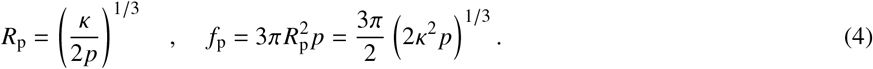

Note that the tube radius scales with the turgor pressure as *R*_p_ ∝ *p*^−1/3^, but the force obeys *f*_p_ ∝ *p*^1/3^. This means a higher turgor pressure results in a narrower tube, but needs larger forces to elongate. In the case of tension-dominant condition, the characteristic tube radius *R_σ_* and force *f_σ_* read (50):

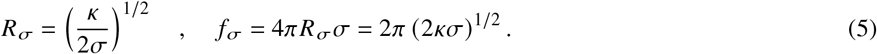

As for endocytosis in yeast cells, *σ* ≈ 0.01 pN/nm (19), *p* ≈ 1 MPa (12, 13) and *κ* ≈ 300*k*_B_T (44). These numbers lead to a rough estimation of *R*_p_ ≈ 8.5 nm, *f*_p_ ≈ 700 pN and *R*_σ_ ≈ 250 nm, *f*_σ_ ≈ 30 pN. The radius of long endocytic invaginations observed experimentally is about 15 nm (14), which is much closer to the estimated *R*_p_ than the estimated *R_σ_*, thus supporting the statement that the turgor pressure but not the membrane tension is the dominant factor that opposes endocytosis in yeast cells. For the rest of the paper, we assume *σ* = 0.002*pR*_p_ such that *R_σ_* = 22*R*_p_ ≫ *R*_p_, and therefore the turgor pressure always dominates over the surface tension in shaping the membrane. We measure the length in units of the characteristic radius *R*_p_ and the force in units of the characteristic force *f*_p_. The pressure is non-dimensionalized with 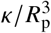 to a constant 0.5. The mechanics of the system is then determined by only a few dimensionless parameters, including the rescaled base radius *R*_b_/*R*_p_, the rescaled spontaneous curvature *c*_0_*R*_p_, as well as the rescaled coating area 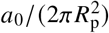 and the rescaled edge sharpness parameter 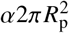 when considering the inhomogeneous model (see Eq. (6)).

### A large base radius lowers the force barrier to pull a membrane tube against turgor pressure

We first consider the case of a membrane at the endocytic site void of any curvature-generating proteins (i.e., *c*_0_*R*_p_ = 0), and study the effect of base radius on the required forces to pull a membrane tube. The effect of forces on the membrane deformation is characterized by the force-height (*f-L*) curve, which in general is non-monotonic (Figure 2a - d). A force barrier *F*_max_ appears at a relatively low height L when the membrane is dome-shaped (Figure 2a - d, inset, labeled by circles). As the membrane is lifted further up, the membrane changes from a dome-shape to an Ω-shape, when a narrow neck appears (signaled by the tangential angle *ψ* = *π*/2 at an intermediate arclength). The force *f* then decreases with L and approaches the elongation force *F*_e_ ≡ lim_*L*→∞_ *f*(*L*), which equals *f*_p_ in the case of a bare membrane as expected by Eq. (4). The existence of a force barrier in the *f-L* curve is similar to that in the tension-dominant condition (50). However, two striking differences should be noted: (i) in the pressure-dominant condition discussed here, a nonzero initiation force *F*_init_ ≡ *f*(*L* = 0) is needed to merely start deforming the membrane, i.e., to lift the membrane just off the cell wall (Figure 2e and f, diamonds), whereas in the tension-dominant condition, *F*_init_ = 0 is independent of *R*_b_ (50); (ii) when pressure dominates, the force barrier *F*_max_ significantly varies with the base radius *R*_b_ (Figure 2e and f, circles), whereas in the tension-dominant condition, *F*_max_ always overshoots 13% relative to the equilibrium force *f_σ_* (50), independent of *R*_b_.

**Figure 2:**
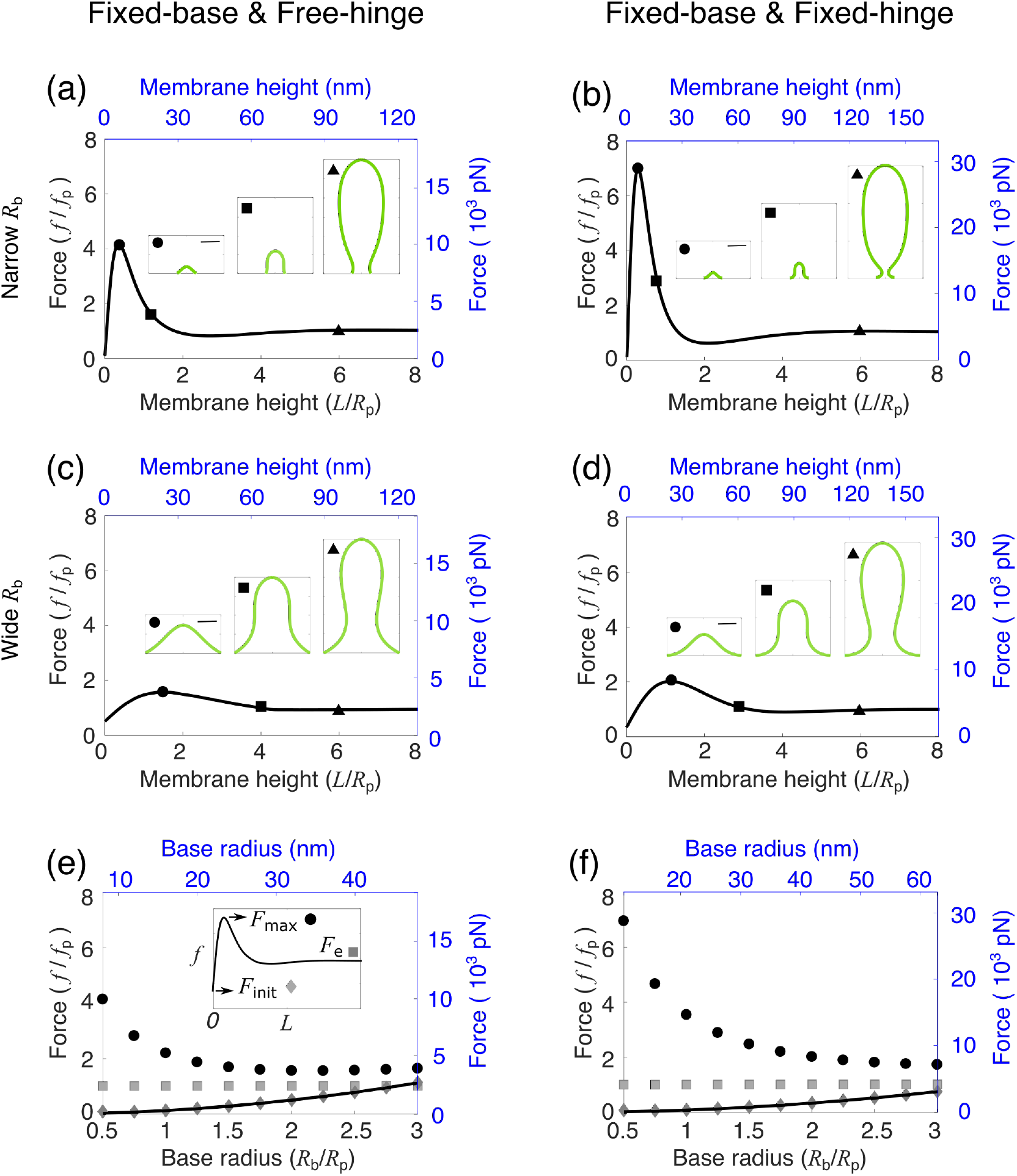
Effect of the base radius *R*_b_ on the membrane shape and force requirement. (a - d) Force-height relationship *f-L* of membrane deformations for a fixed base radius *R*_b_/*R*_p_ = 0.5 in (a, b) and *R*_b_/*R*_p_ = 2 in (c, d), where *R*_p_ is the characteristic tube radius (Eq. 4). The spontaneous curvature *c*_0_*R*_p_ = 0. Insets show membrane shapes at the points indicated by the corresponding symbols on the *f-L* curve. The square indicates the critical shape where the membrane is about to form a neck. The scale bar corresponds to the characteristic tube radius *R*_p_. (e, f) Force barrier *F*_max_ (circle), initiation force *F*_init_ (diamond) and elongation force *F*_e_ (square) for varying base radii *R*_b_. The solid curve represents the analytical solution for *F*_init_. (a-f) In the left column (a, c, e), the free-hinge BC is imposed at the base points *R* = *R*_b_, while in the right column (b, d, f), the fixed-hinge BC is imposed. On the left and bottom axes (black), non-dimensionalized quantities are used, while on the right and top axes (blue), quantities are measured in their physical units. The parameters are listed in Table 2 except *R*_b_ = 8nm in (a) and *R*_b_ = 10.5nm in (b).

When comparing the differences between the *f-L* curves for the two BCs, we notice that: (i) the initiation force *F*_init_ scales with the base radius *R*_b_ as 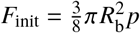 for the free-hinge BC, whereas 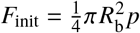 for the fixed-hinge BC (Figure 2e and f, solid curves, see Supporting Material for the derivation); (ii) though the initiation force *F*_init_ is smaller for the fixed-hinge BC than for the free-hinge BC, the opposite trend is observed for the force barrier *F*_max_. The difference in *F*_max_ is more pronounced for smaller base radii. For instance, when *R*_b_ = 0.5*R*_p_, the force barrier *F*_max_ is about 4*f*_p_ for the free-hinge BC, while it is 7 *f*_p_ for the fixed-hinge BC (Figure 2a and b, labeled by circles); (iii) The membrane neck appears at a smaller membrane height for the fixed-hinge BC than for the free-hinge BC. For instance, when *R*_b_ = 2*R*_p_, the neck appears at a height of 3*R*_p_ for the fixed-hinge BC, but 4*R*_p_ for the free-hinge BC (Figure 2c and d, labeled by squares).

When the membrane is pulled up above the height of 6*R*_p_, the force to elongate the tube remains almost unchanged *F*_e_ = *f*_p_, regardless of the BCs and the base radii. However, the shape of the membrane can be quite different for different radii *R*_b_. If *R*_b_ < *R*_p_, the membrane exhibits a balloon-shape with a narrower base than the tubular body (Figure 2a and b, inset, labeled by triangles), whereas when *R*_b_ > *R*_p_, a wider base connected to a narrower body is observed (Figure 2c and d, inset, labeled by triangles), which is more consistent with experimental observations (14).

For both BCs, the force barrier *F*_max_ is significantly reduced with increasing base radius *R*_b_. When the base radius is increased from 0.5*R*_p_ to 3*R*_p_, the force barrier is reduced from 4*f*_p_ to 1.5*f*_p_ for the free-hinge BC, and from 7*f*_p_ to 2*f*_p_ for the fixed-hinge BC (Figure 2e, f, circles). These results suggest that a relatively wide base facilitates CME in yeast cells. With parameters listed in Table 2, the force barrier to pull a membrane tube against a turgor pressure of 1 MPa can be reduced to 3500 pN for the free-hinge BC and 8000 pN for the fixed-hinge BC when the base radius *R*_b_ is greater than 30 nm (Figure 2e, f, circles). For the rest of the paper, we fix the base radius at *R*_b_ = 2*R*_p_ and study the other factors that influence the membrane shape and the force to pull a membrane tube.

**Table 2:** Fitting parameters to compare with experimental data

| Symbols | Physical meaning | Values for the free-hinge BC | Values for the fixed-hinge BC |
| --- | --- | --- | --- |
| $p$ | Turgor pressure | 1 MPa | 1 MPa |
| $R_p$ | Characteristic tube radius | 16 nm | 21 nm |
| $c_0$ | Spontaneous curvature of the membrane induced by protein coat | $0.063 \text{ nm}^{-1}$ | $0.048 \text{ nm}^{-1}$ |
| $a_0$ | Coating area of proteins | $1609 \text{ nm}^2$ | $2771 \text{ nm}^2$ |
| $R_b$ | Base radius of the membrane patch | 32 nm | 42 nm |
| $\sigma$ | Surface tension at the base | $0.032 \text{ pN/nm}$ | $0.042 \text{ pN/nm}$ |
| $\alpha$ | Control parameter for the sharpness of the coating edge | $0.006 \text{ nm}^{-2}$ | $0.004 \text{ nm}^{-2}$ |

### The ability of a fully-covered protein coat to reduce the force barrier and the initiation force depends on boundary conditions

In this section, we consider the effect of a uniform coat of curvature-generating proteins on membrane deformations. The ability of curvature-generating proteins to induce membrane curvature is characterized by the spontaneous curvature *c*_0_ in the model. When the spontaneous curvature *c*_0_ is small, e.g., *c*_0_*R*_p_ = 0.2, the *f-L* curves show similar trends as a fully uncoated coated membrane. However, a new branch of solutions with negative forces emerges (Figure 3a and b, dashed line). On this branch, the membrane exhibits a highly curved Ω-shape, and has part of the shape lying below the plane *z* = 0. The membrane therefore may interact with the cell wall. This interaction is not considered in our model. The branch terminates at a limiting shape of a closed spherical vesicle budding off from the base (Figure 3a and b, inset, labeled by stars). The free energy of the membrane on this negative-force branch is significantly higher than that on the positive-force branch (Figure S1 in the Supporting Material), thus being energetically unfavorable. Hereafter, the free energy refers to Eq. (1) excluding the contribution – *fL* from the external pulling force.

**Figure 3:**
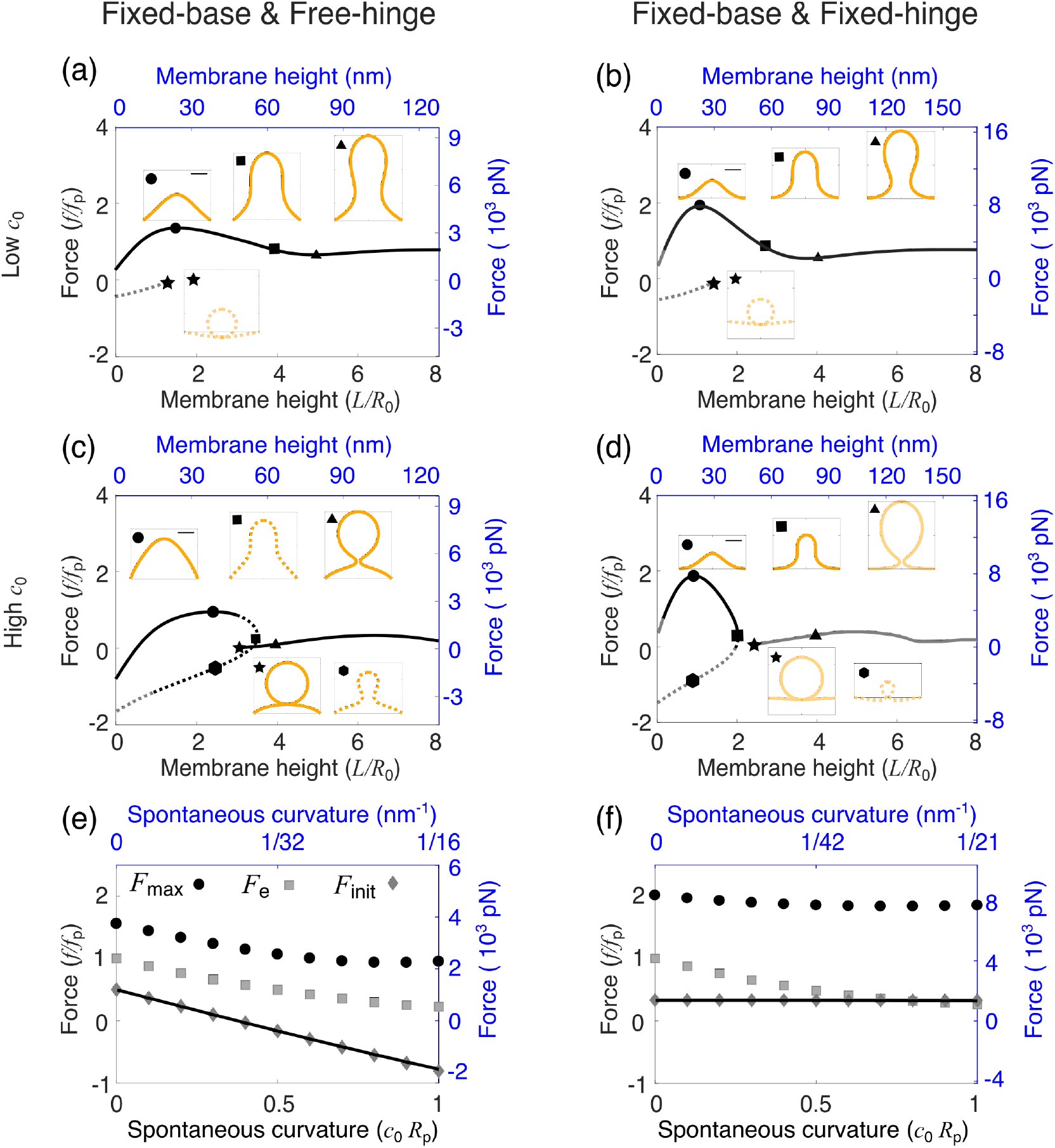
Effect of the spontaneous curvature *c*_0_ on membrane shape and force requirement for a fully coated membrane. (a - d) Force-height (*f-L*) relationship of membrane deformations for a fixed spontaneous curvature *c*_0_*R*_p_ = 0.2 in (a, b) and *c*_0_*R*_p_ = 1 in (c, d). Insets show membrane shapes at the points indicated by the corresponding symbols on the *f-L* curve. The square indicates the critical shape where the membrane is about to form a neck. The scale bar corresponds to *R*_p_. In (a - d), the solid line indicates shapes of the lowest free energy and the dashed line indicates shapes of relatively high free energy. The dark color indicates membrane shapes that are all above *z* = 0, and the gray color indicates shapes that have parts below *z* = 0. (e, f) Force barrier *F*_max_ (circle), initiation force *F*_init_ (diamond) and elongation force *F*_e_ (square) for varying *c*_0_. The solid curve represents the analytical solution for *F*_init_. (a - f) In the left column (a, c, e), the free-hinge BC is imposed at the base points *R*_b_ = 2*R*_p_, while in the right column (b, d, f), the fixed-hinge BC is imposed. On the left and bottom axes (black), non-dimensionalized quantities are used, while on the right and top axes (blue), quantities are measured in their physical units. The parameters are listed in Table 2.

When the spontaneous curvature *c*_0_ is large, e.g., *c*_0_*R*_p_ = 1, the *f-L* curve breaks into two branches, each branch only covering part of the membrane height (Figure 3c and d). In the small-*L* branch, one *L* has two corresponding forces *f*. The larger *f* corresponds to a solution with a dome shape (Figure 3c and d, inset, labeled by circles), while the smaller *f* corresponds to a solution with an Ω-shape (Figure 3c and d, inset, labeled by hexagons). The dome-shape has lower free energy than the Ω-shape for the same membrane height *L*, and therefore is energetically more stable (Figure S1, c and d). The large-*L* branch starts from a point at which the force *f* is slightly above zero, and the shape of the membrane is shown as a vesicle budding off from the base (Figure 3c and d, inset, labeled by stars). This shape has the lowest free energy in the large-*L* branch, which implies that if a long tube is pulled up and maintained by a force, when the force is gradually released, the tube retracts and a vesicle spontaneously forms and detaches from the base of the membrane.

For a fully coated membrane, increasing the spontaneous curvature *c*_0_ is able to reduce the elongation force *F*_e_. With increasing *c*_0_*R*_p_ from 0 to 1, *F*_e_ shows exactly the same dependence on *c*_0_ for both BCs and drops from *f*_p_ to about 0.2*f*_p_ (Figure 3e and f, squares). However, The impact of the spontaneous curvature *c*_0_ on the initiation force *F*_init_ and the force barrier *F*_max_ shows qualitative differences between the two BCs: (i) under the free-hinge BC, the initiation force *F*_init_ drops down with increasing *c*_0_ and becomes negative for *c*_0_*R*_p_ > 0.4 (Figure 3e, diamonds and solid line). This negative *F*_init_ implies that the membrane is spontaneously bent off the cell wall without external forces. By contrast, under the fixed-hinge BC, the initiation force *F*_init_ remains positive and almost constant (Figure 3f, diamonds and solid line); (ii) the force barrier *F*_max_ noticeably decreases from 1.5*f*_p_ to *f*_p_ with increasing *c*_0_ under the free-hinge BC, while *F*_max_ remains almost constant at 2*f*_p_ under the fixed-hinge BC (Figure 3e and f, circles).

In biological terms, our results suggest that for a membrane fully coated with curvature-generating proteins, the protein coat might significantly reduce the forces to start deforming the membrane if the membrane angle at the base is free to rotate. However, the protein coat has little impact on the forces if the membrane angle is fixed to zero. With the parameters listed in Table 2, the force barrier to pull a membrane tube for the fixed-hinge BC can be reduced from 3500 pN for a fully uncoated membrane to 2500 pN for a fully-coated membrane (Figure 3e, circles), but the force barrier is kept at 8000pN for the fixed-hinge BC, regardless of the spontaneous curvature (Figure 3f, circles).

### Partially coating the membrane with curvature-generating proteins can reduce the initiation force and the force barrier but not the elongation force

In this section, we study the inhomogenenous model where the membrane is coated with curvature-generating proteins only around the tip, thus mimicking the distribution of clathrin and other adaptor proteins. The spontaneous curvature in our model spatially varies as

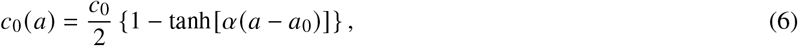

where *a*(*s*) is the surface area calculated from the tip to the position of arclength *s*. The parameter *α* controls the sharpness of the coating edge. The coating area of proteins is denoted by *a*_0_, and these proteins induce a spontaneous curvature of *c*_0_ in the coated region of the membrane. This form of spontaneous curvature has been used in many previous studies (16, 17, 19, 20).

We first vary the coating area a_0_ while fixing the spontaneous curvature at *c*_0_*R*_p_ = 1. When a_0_ is small, the *f-L* curves are non-monotonic with a single force barrier 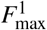 at a low membrane height, similar to that of a bare membrane (data not shown). However, when *a*_0_ is above a critical value, a second force barrier 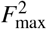 emerges on the *f-L* curve at a higher membrane height where the membrane exhibits an Ω-shape (Figure 4a and b, inset, labeled by triangles). For 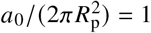, the protein coat forms a hemispherical cap when the membrane is pulled up into a tubular shape (Figure 4a and b, inset, labeled by triangles). The initiation forces are negative for both BCs and the zero force *f* = 0 intersects with the *f-L* curve at a positive membrane height (Figure 4a and b, inset, labeled by circles). For very large coating area 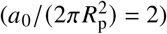, the membrane is almost fully coated with proteins when the membrane is flat (Figure 4c, inset, labeled by circles). The *f-L* curve is broken into two branches, each branch only covering part of the membrane height (Figure 4c and d), similar to the *f-L* curve of a fully coated membrane. The two branches might overlap in some intermediate membrane heights. For the free-hinge BC, the zero force *f* = 0 intersects with the *f-L* curve at three points, two of them lying on the small-*L* branch and the third one on the large-*L* branch (Figure 4c, inset, labeled by circles and squares in the small-*L* branch and triangles in the large-*L* branch). The two points on the small-*L* branch correspond to a dome-shape of low free energy and a tubular shape of high free energy (Figure S2, c, circles and squares). Therefore, in the absence of forces, the membrane adopts a dome-shape, spontaneously curved off from the cell wall. The one point on the large-*L* branch corresponds to a highly curved Ω-shape with a narrow neck (Figure 4c, inset, labeled by triangles), which is the final shape of a long membrane tube when it retracts upon force release. The large-*L* branch starts with a limiting membrane shape that is a closed vesicle budding off from the base (Figure 4c, inset, labeled by stars). In contrast with the fully coated membrane, the force at this point is negative, which means that a downward force is further needed to push the membrane into a budding vesicle when the membrane tube retracts. Under the fixed-hinge BC, the *f-L* curve for 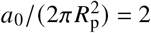 shows similar features with that of the free-hinge BC, except that the dome-shaped solution at *f =* 0 does not exist (Figure 4d). This is because the initiation force *F*_init_ is positive and the membrane cannot be spontaneously curved off from the cell wall.

**Figure 4:**
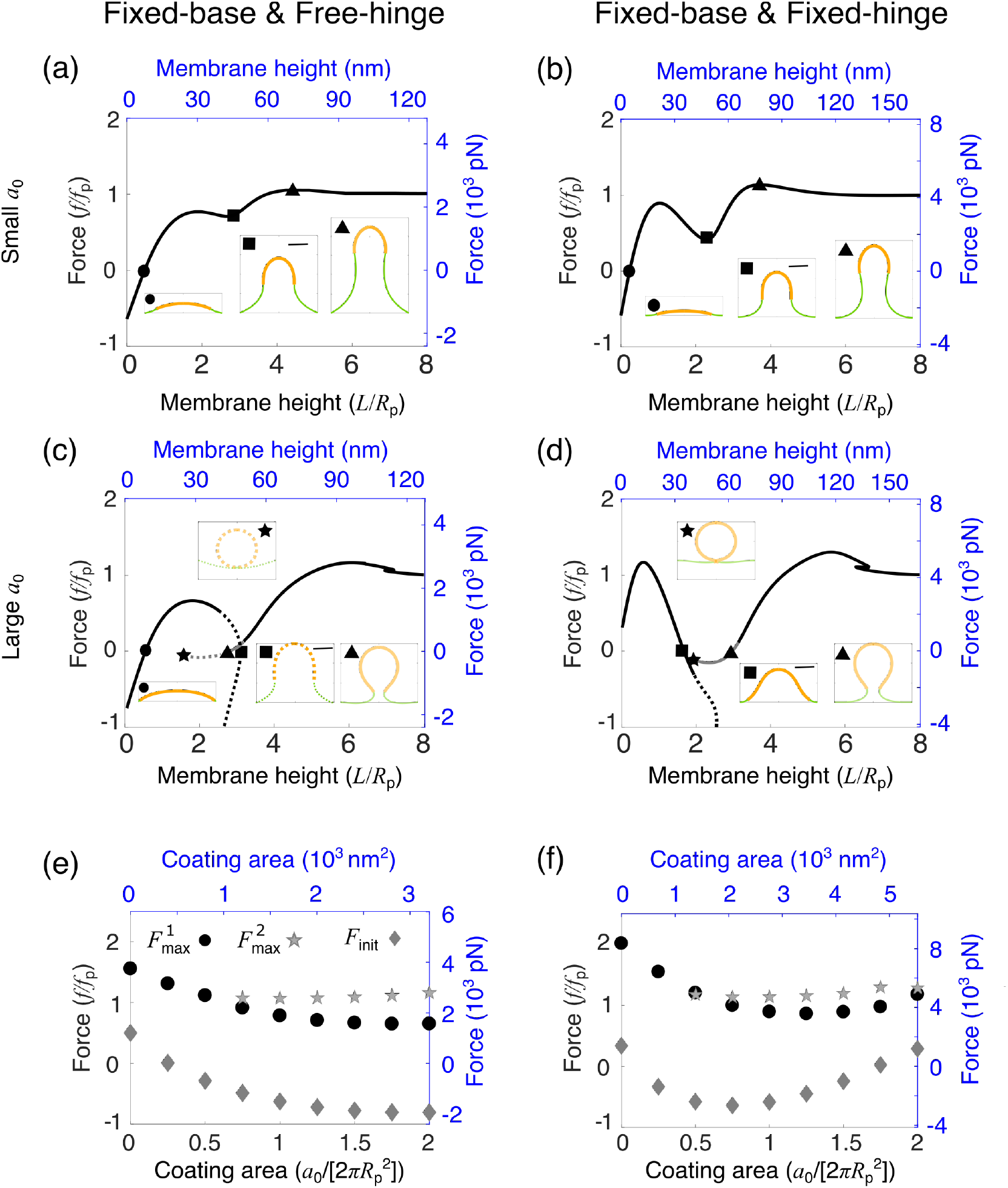
Effect of the coating area *a*_0_ of curvature-generating proteins on membrane shape and force requirement for a partially coated membrane. (a - d) Force-height (*f-L*) relationship of membrane deformations for a fixed coating area 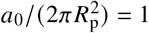 in (a, b) and 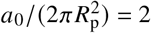 in (c, d). Insets show membrane shapes at the points indicated by the corresponding symbols on the *f-L* curve. The orange part represents the area of the membrane coated with proteins and the green part represents the bare membrane. The scale bar corresponds to *R*_p_. In (a - d), the solid line indicates shapes of the lowest free energy and the dashed line indicates shapes of relatively high free energy. The dark color indicates membrane shapes that are all above *z* = 0, and the gray color indicates shapes that have parts below *z* = 0. (e, f) Low-height force barrier 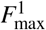 (circle), high-height force barrier 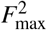 (star) and initiation force *F*_init_ (diamond) for varying *a*_0_. (a - f) In the left column (a, c, e), the free-hinge BC is imposed at the base points *R*_b_ = 2*R*_p_, while in the right column (b, d, f), the fixed-hinge BC is imposed. On the left and bottom axes (black), non-dimensionalized quantities are used, while on the right and top axes (blue), quantities are measured in their physical units. The parameters are listed in Table 2.

Despite some common features in the *f-L* curves for both BCs, differences also exist: (i) under the free-hinge BC, the initiation force *F*_init_ decreases and remains negative with increasing *a*_0_, whereas under the fixed-hinge BC, *F*_init_ is negative for intermediate values of *a*_0_, and becomes positive for larger *a*_0_ (Figure 4e and f, diamonds); (ii) a similar difference is also observed for the low-height force barrier 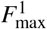, which is monotonically decreasing with *a*_0_ under the free-hinge BC, whereas it is non-monotonic under the fixed-hinge BC (Figure 4e and f, circles).

For a partially coated membrane, the low-height force barrier 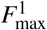 can be significantly reduced to below *f*_p_ for some coating areas (Figure 4e and f, circles), whereas the high-height force barrier 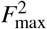 increases with *a*_0_ and remains above *f*_p_ (Figure 4e and f, stars). This is because the force barrier 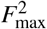 must be greater than the elongation force *F*_e_, which equals *f*_p_ for both BCs and any coating areas. This trade off between the two force barriers implies there is an optimum coating area that minimizes the overall force barrier. With the parameters listed in Table 2, the optimum coating area is about 1200 nm^2^ for the free-hinge BC and 2000 nm^2^ for the fixed-hinge BC. The minimum force barrier is about 2500 pN for the free-hinge BC, and about 4000pN for the fixed-hinge BC. Compared with the force barrier of 8000pN for a fully coated membrane under the fixed-hinge BC, partially coating the membrane significantly reduces the force barrier.

### Increasing the spontaneous curvature of partially coated membrane leads to a sharp transition of the membrane shape

In this section, we vary the spontaneous curvature *c*_0_ while fixing the coating area 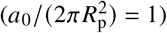 to study how *c*_0_ influences the *f-L* curves for a partially coated membrane. Upon gradually increasing *c*_0_, the *f-L* curve shows similar trends to what we observed when increasing the coating area. Above a critical value of *c*_0_, a high-height force barrier 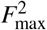 appears on the *f-L* curve in addition to the low-height force barrier 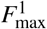 (Figure 5a and b). Further increasing the spontaneous curvature *c*_0_ splits the *f-L* curve into two branches, a small-*L* branch and a large-*L* branch (Figure 5c and d). A striking new feature is that when *c*_0_*R*_p_ = 2, the force for the entire small-*L* branch falls below zero (Figure 5c and d). The zero force *f* = 0 intersects with the *f-L* curve on the long-*L* branch at only one point, which corresponds to a highly curved Ω-shape (Figure 5c and d, inset, labeled by squares). This shape has the lowest free energy (Figure S3, c and d, labeled by squares), which implies that even in the absence of forces, increasing the spontaneous curvature *c*_0_ can lead to a transition of the membrane from the dome-shape in the small-*L* branch to the Ω-shape in the large-*L* branch. The membrane height has a sharp increase during this transition.

**Figure 5:**
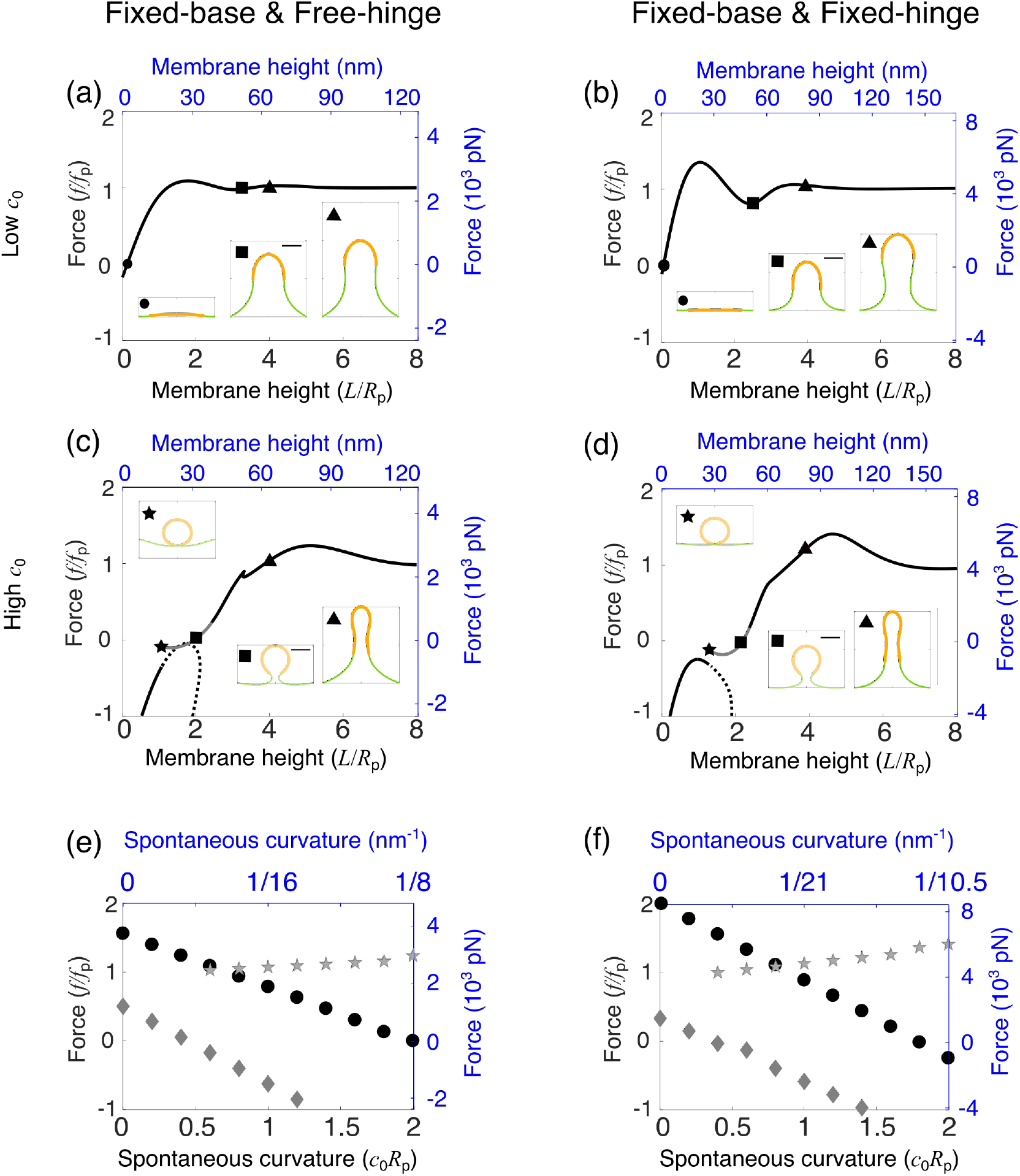
Effect of the spontaneous curvature *c*_0_ of curvature-generating proteins on membrane shape and force requirement for a partially coated membrane. (a - d) Force-height (*f-L*) relationship of membrane deformations for a fixed spontaneous curvature *c*_0_*R*_p_ = 0.6 in (a, b) and *c*_0_*R*_p_ = 2 in (c, d). Insets show membrane shapes at the points indicated by the corresponding symbols on the *f-L* curve. The orange part represents the area of the membrane coated with proteins and the green part represents the bare membrane. The scale bar corresponds to *R*_p_. In (a - d), the solid line indicates shapes of the lowest free energy and the dashed line indicates shapes of relatively high free energy. The dark color indicates membrane shapes that are all above *z* = 0, and the gray color indicates shapes that have parts below *z* = 0. (e, f) Low-height force barrier 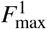 (circle), high-height force barrier 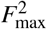 (star) and initiation force *F*_init_ (diamond) for varying *c*_0_. (a - f) In the left column (a, c, e), the free-hinge BC is imposed at the base points *R*_b_ = 2*R*_p_, while in the right column (b, d, f), the fixed-hinge BC is imposed. On the left and bottom axes (black), non-dimensionalized quantities are used, while on the right and top axes (blue), quantities are measured in their physical units. The parameters are listed in Table 2.

The spontaneous curvature *c*_0_ not only influences the forces but also the morphology of the clathrin coat. When *c*_0_*R*_p_ = 2, the clathrin coat tends to bend the membrane to a narrow radius of ≈ 0.5*R*_p_ and the coated area exhibits a pearl-like structure when elongated (Figure 5c and d, triangles). However, for *c*_0_*R*_p_ = 1, the clathrin coat maintains a roughly hemispherical cap (Figure 4a and b, triangles).

Both the low-height force barrier 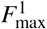 and the initiation force *F*_init_ linearly decrease with increasing *c*_0_ (Figure 5e and f, circles and diamonds), and they become negative when *c*_0_ is beyond a critical value. By contrast, the high-height force barrier 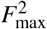 linearly increases with *c*_0_ (Figure 5e and f, stars), and remains above *f*_p_. The optimum spontaneous curvature, which has the minimum force barrier, is about 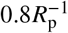 for both BCs. The corresponding force barrier is as much as *f*_p_, which is the lowest force barrier one can achieve by partially coating the membrane with curvature-generating proteins. With the parameters listed in Table 2, the optimum spontaneous curvature corresponds to a preferred radius of about 40 nm for the free-hinge BC and 50 nm for the fixed-hinge BC. The force barrier for the free-hinge BC is about 2500pN, and for the fixed-hinge BC is about 4000pN.

### Our theory agrees well with experiments

The shapes of endocytic invaginations in budding yeast have been imaged with electron tomography (14). These shapes typically do not have perfect axisymmetry assumed in our model (Figure 6a and b). However, from these images one can numerically fit the membrane shape and extract geometric features of the shape, which typically include the tip radius *R*_t_, the tip-neck distance *D*_t_ and the membrane height *L* (14). The tip radius *R*_t_ is defined as the reciprocal of the meridian curvature 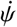 averaged over an arc that extends over 15 nm from the endocytic invagination tip. The tip-neck distance *D*_t_ is defined as the distance from the center of the neck to the most distant profile point from the neck. The membrane height *L* is defined as the maximum height of the fitted profile above the base. The experimental datasets *R*_t_ v.s. *L* and *D*_t_ v.s. *L* contain the shape information of the endocytic invagination across different stages of CME. We use the two datasets as the fitting data to compare our theory with experiments. The fitting procedure is elaborated in the Appendix, where we use the characteristic radius *R*_p_ as the single parameter to fit the data. We find the optimum 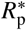 that minimizes the fitting error for the two datasets. For the free-hinge BC, the optimum 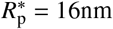, and for the fixed-hinge BC 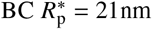 (Figure S4). The fitting errors at the optimum 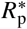 are comparable for the two BCs, and we cannot distinguish which BC fits the experimental data better (Figure S4).

**Figure 6:**
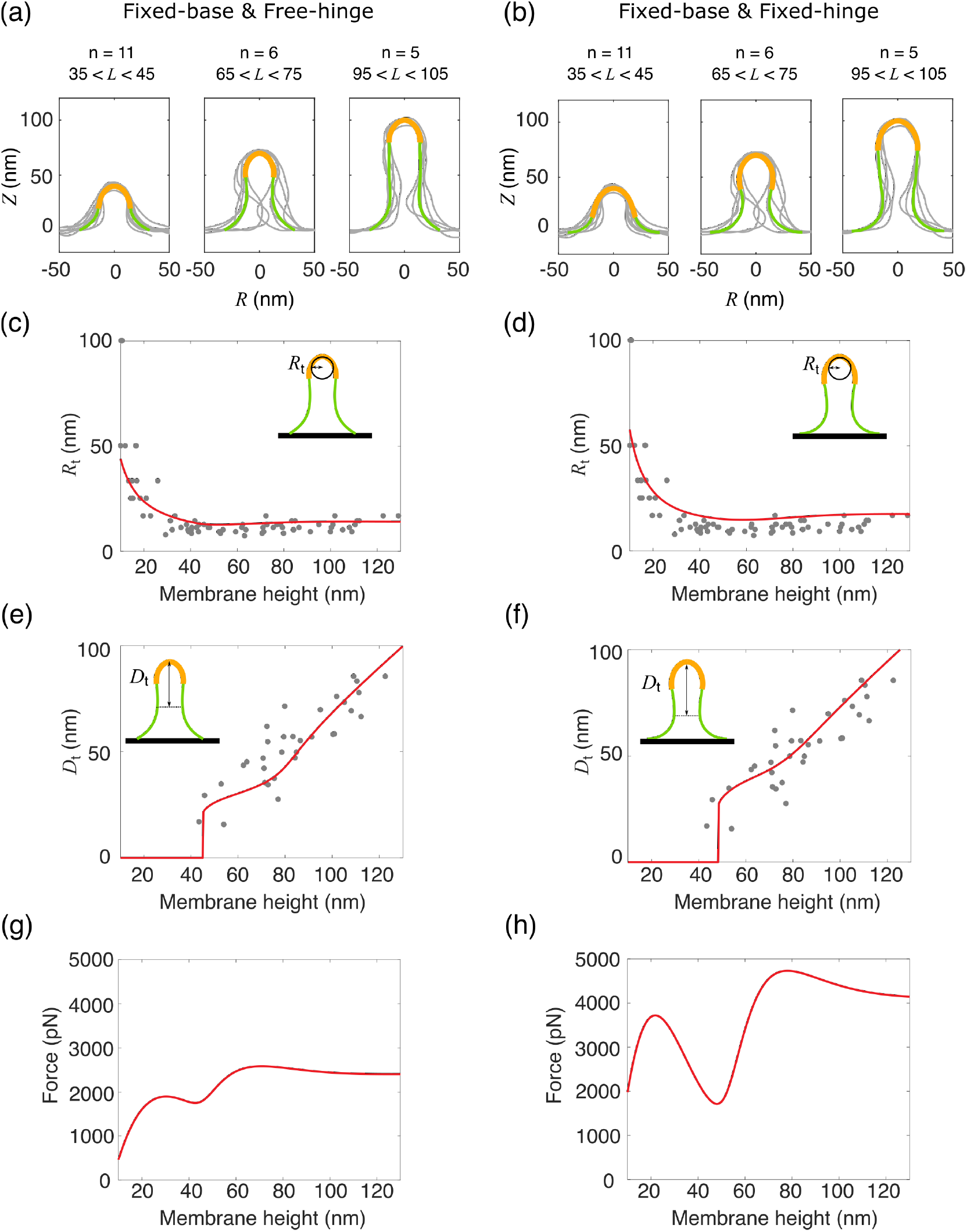
Comparison between our theory and experiments. (a, b) Membrane shapes obtained by electron to-mography are grouped according to their heights and overlaid at the tip. The data come from the database https://www.embl.de/download/briggs/endocytosis.html maintained by the authors of Ref. (14). The membrane shapes obtained by our model are represented by solid curves. The orange part represents the area of the membrane coated with proteins and the green part represents the bare membrane. (c, d) Comparison of the tip radius *R*_t_ between obtained with our theory (line) and measured experimentally (dots). (e, f) Comparison of the neck to tip distance *D*_t_ between obtained with our theory (line) and measured experimentally (dots). (g, h) Prediction of the force-height (?-L) relationship from our theory using the parameters listed in Table 2 which fit the experimental shapes in (a, b). (a-h) In the left column (a, c, e, g), the free-hinge BC is imposed at the base points *R*_b_ = 32 nm, while in the right column (b, d, f, h), the fixed-hinge BC is imposed at the base points *R*_b_ = 42 nm.

Using the optimum 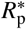, our calculated membrane shapes agree well with the experimental profile, particularly in the early stage when the membrane height is low (Figure 6a and b). For membrane shapes that are higher than 65 nm, experimental membrane shapes are typically asymmetric and exhibit a narrower neck than the calculated ones, probably due to the presence of other membrane proteins that arrive later during CME and impose a cylindrical curvature at the neck of the invagination (e.g. amphiphysins). These effects are not considered in our model.

As for the geometric features, experimental data shows that the tip radius *R*_t_ drops from 50 – 100nm to 15nm as the membrane height increases. Our theory matches the trend of the experimental data, particularly for the part where *R*_t_ < 40 nm (Figure 6c and d). The fitting for the tip radius with the free-hinge BC is slightly better than that with the fixed-hinge BC. For the tip-neck distance *D*_t_, our theory predicts that *D*_t_ grows slowly with the membrane height L when L is less than 65 nm. Beyond this point, *D*_t_ scales almost linearly with *L* with a larger slope than the initial phase. This theoretical prediction again matches well with the experimental data (Figure 6e and f). The fitting for the tip-neck distance with the fixed-hinge BC is slightly better than that with the fixed-hinge BC.

We stress that the different optimum 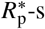 for the two BCs result in a large difference in the magnitude of forces in the *f-L* curve (Figure 6g and h). This is because the unit of the force is the characteristic force *f*_p_ which scales with the characteristic radius *R*_p_ as 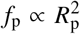. As a result, the force barrier is about 2500 pN for the free-hinge, while it is about 4000 pN for the fixed-hinge.

### Fixed base v.s. freely moving base

We have focused on BCs where the base radius of the membrane is fixed. For a membrane fully coated with curvature-generating proteins, the initiation force *F*_init_ either decreases with the intrinsic curvature *c*_0_ under the free-hinge BC, or is independent of *c*_0_ under the fixed-hinge BC (Figure 3e and f, diamonds and solid lines). A previous work (18) studied a similar homogeneous model but used the free-base and fixed-hinge BC (BC4 in TABLE 1). This BC led to the surprising conclusion that the initiation force *F*_init_ of a fully coated membrane is proportional to the spontaneous curvature *c*_0_, which implies that increasing the spontaneous curvature *c*_0_ hinders CME because it raises the force required to lift the membrane off the cell wall. In addition, as a result of the freely moving base, the model predicted that the base radius *R*_b_ approaches zero when the membrane height is low. This result is inconsistent with experimental observations that the base radius of membrane invaginations remains roughly the same during the entire course of CME, from shallow invaginations to long tubes (Figure 6a and b). Therefore the experimental data supports our assumption that the base of the membrane is maintained at a fixed radius by endocytic proteins or by attachment to the cell wall. A recent systematic study of proteins involved in endocytosis by super-resolution microscopy revealed that many proteins are organized in concentric rings around the clathrin coat (28). These proteins may serve as anchors and may fix the base radius of the endocytic membrane. In addition, the actin network around the endocytic invagination can also impose constrains on the extent the membrane base can spread. Furthermore, we fix the surface tension at the fixed base radius. It implies the assumption that the lipids can flow past any structures that fix the membrane to the cell wall.

The different dependence of the initiation force *F*_init_ on *c*_0_ between the fixed-base BC and the free-base BC can be clarified with a simple example. Since *F*_init_ is only related to the early stage of CME when the membrane is almost flat, we approximate the dome-shaped membrane as a spherical cap and calculate its free energy *E*(*R*; *c*_0_, *R*_b_) as a function of the sphere radius R for different spontaneous curvatures *c*_0_ and base radii *R*_b_ (Figure 7). For the fixed-base BC, the base radius *R*_b_ is a constant. When *c*_0_ is small, *E*(*R*; *c*_0_, *R*_b_) decreases monotonically with R and has its minimum at *R* = ∞, which implies that a flat shape is more favorable than a curved one (Figure 7a). When *c*_0_ becomes large, E(R; *c*_0_, *R*_b_) has a nontrivial minimum at a finite radius *R* (Figure 7b,*R*_b_ = 2*R*_p_), which implies that the membrane spontaneously bends into a curved shape. However, for the free-base BC assumed in the work of (18), the base radius *R*_b_ becomes a free parameter and the free energy *E*(*R*, *R*_b_; *c*_0_) is a function of both R and *R*_b_. No matter how large *c*_0_ is, the energy *E*(*R*, *R*_b_; *c*_0_) always admits a trivial minimum at *R*_b_ = 0, which represents a solution without any deformation (Figure 7a and b, *R*_b_ = 0*R*_p_). If a force *f* is applied, a nontrival minimum of the total free energy *F*(*R*, *R*_b_; *f*, *c*_0_) = *E*(*R*, *R*_b_; *c*_0_) – *fL*(*R*, *R*_b_) may exist for a positive force *f* (Figure S5). However, the base radius for this nontrivial minimum is unrealistically narrow (~ 0.02nm, see Supporting Material), therefore a freely moving base is probably not a proper BC to model CME in yeast.

**Figure 7:**
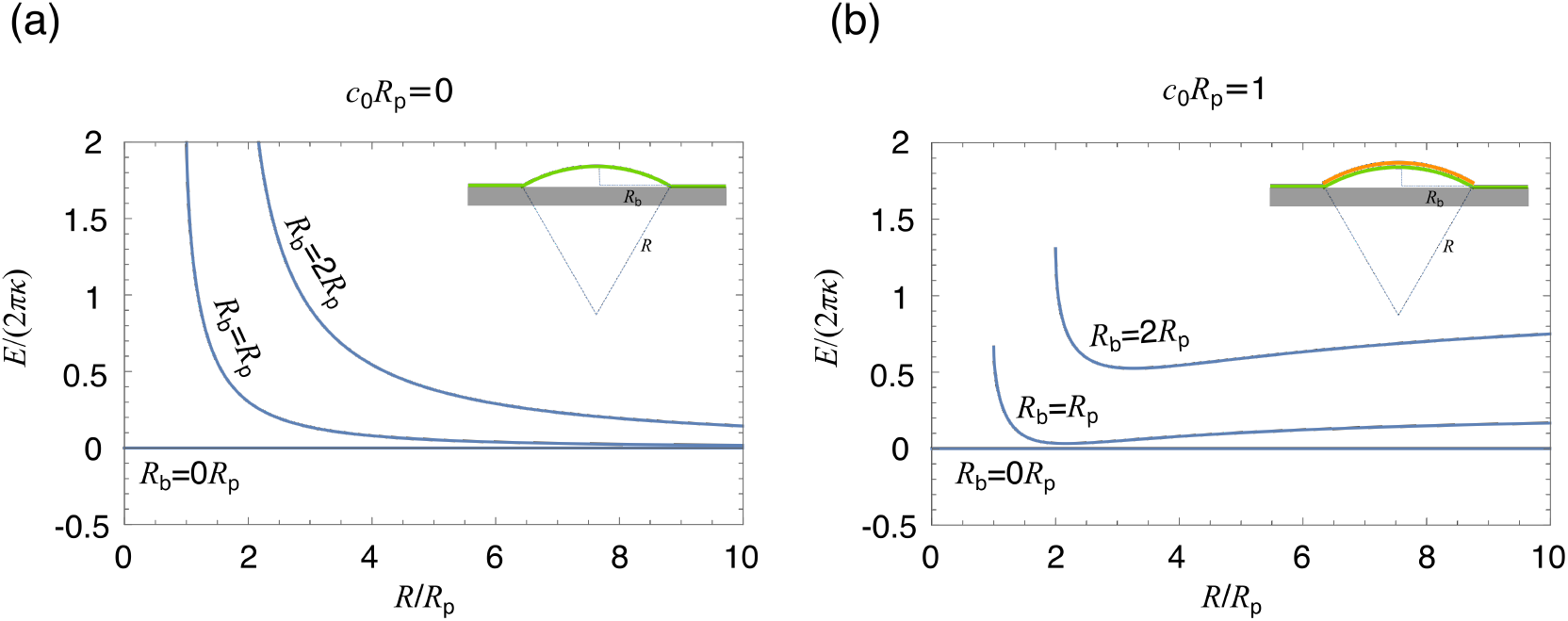
Free energy of membrane deformations under spherical cap approximation. (a, b) Free energy of the membrane as a function of the sphere radius R for *c*_0_*R*_p_ = 0 in (a) and *c*_0_*R*_p_ = 1 in (b). For different base radii *R*_b_, the range of *R* is [*R*_b_, ∞], where *R* = *R*_b_ corresponds to a hemi-spherical cap and *R* = ∞ corresponds to a flat shape.

## DISCUSSION

### Free-hinge v.s. fixed-hinge

Our analysis of the experimental data favors the BC with fixed base radius over that with freely moving base. However, we cannot directly distinguish whether the angle of the membrane at the base is free to rotate (free-hinge) or fixed to zero (fixed-hinge), since both BCs show good agreements with the experimental data (Figure 6a-f). Under the free-hinge BC, the membrane shape has a kink at the base points. We stress that this discontinuity in the membrane angle is physically and biologically plausible. First, for a membrane fully-coated with curvature generating proteins, the membrane’s spontaneous curvature can change abruptly at the base points and such discontinuity of the mechanical properties of the membrane will result in a kink. Second, for a partially-coated membrane whose mechanical properties smoothly change across the base points, the kink can be induced by external factors. Though it is hypothetical, early arriving endocytic proteins, such as myosin-I motors and BAR-domain proteins Syp1p, Cdc15p and Bzz1p form a ring-like structure around the clathrin-coated pit (28). The microscopic interactions between the ring, the membrane and the cell wall determine the exact BCs. At the macroscopic level, the phenomenological method of membrane mechanics used in this paper allows the presence of a kink as long as the underlying microscopic interactions permit. The free-hinge BC is only one of the many possible BCs that form a kink. Even for the fixed-hinge BC, the fixed angle is not necessarily zero but determined by the microscopic interactions. When tuning the membrane angle at the base for the fixed-hinge BC, we notice that the force barrier to pull a bare membrane into a tube can be reduced by increasing the base angle (Figure S6).

Our calculations assume a single type of BCs for the entire stage of CME. We have shown that the free-hinge BC and the fixed-hinge BC might lead to dramatically different *f-L* curves. These results suggest a new way to regulate CME by tuning the BCs. By changing the BC from the fixed-hinge to the free-hinge, the force barrier is typically reduced. If at early stage the BC is fixed-hinge, switching to free-hinge permits the accumulated force to drive the transformation of the membrane from a dimple shape to a tubular shape, providing the force is larger than the force barrier determined by the free-hinge BC but smaller than the force barrier determined by the fixed-hinge BC.

### Homogeneous model v.s. inhomogeneous model

We have studied not only the homogeneous model, i.e., a fully coated (or fully uncoated) membrane, but also the inhomogenous model, i.e., a partially-coated membrane. Comparing the two models, we noted the following differences: (i) In the inhomogeneous model, two force barriers in the *f-L* curve emerge as the spontaneous curvature *c*_0_ increases, and the low-height force barrier can be significantly reduced, even to values below zero, with increasing *c*_0_ (Figure 5e and f, circles). However, in the homogeneous model, there is only one force barrier, which can be hardly reduced with increasing *c*_0_, especially in the fixed-hinge BC (Figure 3e and f, circles). (ii) The elongation force *F*_e_ can be reduced with *c*_0_ in the homogeneous model (Figure 3e and f, squares), while in the inhomogeneous model it remains at a constant value of *f*_p_ regardless of BCs and parameter values of *α*_0_ and *c*_0_. These differences suggest that a partially-coated membrane can be spontaneously lifted up to a significant height via the curvature-generating protein coat, while it is impossible to do so when the membrane is fully coated.

### Actin polymerization alone is insufficient to overcome the force barrier for CME in yeast cells even with the help of proteins that induce membrane curvature

One of the key questions we aimed to address in this paper is how much force is needed to pull a membrane tube against high turgor pressure during CME. We have assumed a turgor pressure of 1 MPa and estimated that the force barrier is about 2500 pN for the free-hinge BC, but 4000 pN for the fixed-hinge BC (Figure 6g and h). In this calculation, we have assumed a point force acting on the membrane, which is a good approximation if the forces produced by actin filaments are concentrated near the tip of the membrane since the point force is the limit of a concentrated force distribution. We expect the point force is the most efficient way to deform a flat membrane into a tubular shape since it minimizes the total amount of force necessary to deform the membrane. Indeed, let’s consider a concentrated force distribution acting on the membrane such that the normal stress is larger than the turgor pressure at the stress-applied area. The stress is able to overcome the turgor pressure, therefore pulls the membrane up locally, and the stress-free parts of the membrane are raised up correspondingly. If the same amount of force is distributed on a larger area, the resulting stress is reduced and might be smaller everywhere on the membrane than the turgor pressure, and therefore could not pull the membrane up. As an attempt to test this idea, we calculated the *f-L* curve assuming the forces are distributed within an area of *a*_f_ near the membrane tip and pointing in the normal direction. In agreement with our expectation, when the forces are distributed on a larger area *a*_f_, a larger force *f* is needed to pull up the membrane and the increase in force magnitude can be more than 2-fold (Figure S8). Based on this argument, we expect our results provide a lower bound for the force barrier. However, even 2500pN is still beyond the force (< 200pN) that can be generated by polymerization alone of 150 – 200 actin filaments at the endocytic site (46, 51), given that the measured polymerization force for a single filament is only 1pN and the force generated by a bundle of filaments is usually smaller than the sum of each individual ones (52). Investigating non-polymerization based force production by the actin machinery will be our future work. A possible way is to release the elastic energy stored in geometrically frustrated crosslinkers, such as fimbrin (53, 54). There are some important factors about CME in yeast cells we do not consider in our model. The clathrin coat in general is stiffer than the plasma membrane (44). Therefore, the clathrin-coated part of the membrane should have a larger bending rigidity κ than the uncoated part. There are also BAR proteins that bind to the side of the membrane invagination and they impose an anisotropic curvature on the membrane. Investigating these missing factors will help us provide a more accurate estimation of the force requirement to pull the membrane up against turgor pressure for CME in yeast cells.

## CONCLUSION

We have studied membrane deformations driven by a point force and by curvature-generating proteins in the presence of a high turgor pressure. A significant amount of force is required to deform the membrane as a result of the high turgor pressure. We have investigated possible ways to reduce the force requirement. This includes fully or partially coating the membrane with curvature-generating proteins and letting the membrane angle at the base freely rotate. By comparing with experimental data, we have shown that the BC with a fixed base radius is more appropriate than the freely moving base in describing membrane invaginations at the endocytic sites. The minimum force barrier predicted by our theory is about 2500 pN.

## APPENDIX

### Derivation of the membrane shape equations

The membrane shape is parameterized with its meridional coordinates [*R*(*s*), *z*(*s*)], which are related to the tangent angle *ψ*(*s*) via the goemetrical relation:

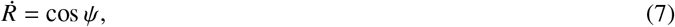

and

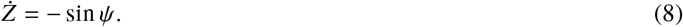

In order to obtain the Euler-Lagrange equation associated with the free energy Eq. (1), we express 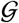 in Eq. (2) explicitly as

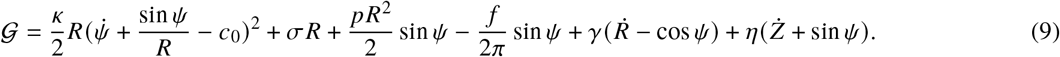

Here we introduce the rescaled Lagrangian multiplier 2*πγ*(*s*) and 2*πņ*(*s*) to impose the geometric constraints set by Eqs. (7) and (8). The variation of the functional *G* in Eq. (2) reads

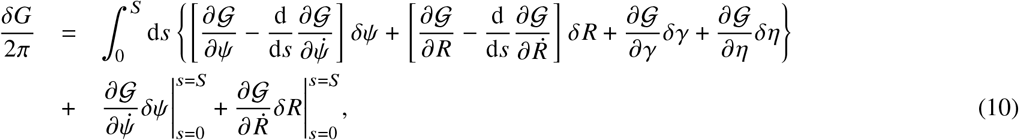

which contains both the bulk terms (first line) and the boundary terms (second line). The Euler-Lagrange equations can be obtained by the vanishing bulk terms, which are reduced to

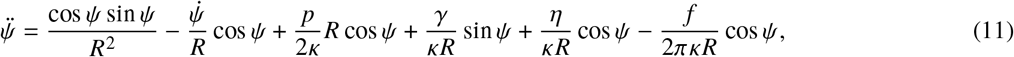

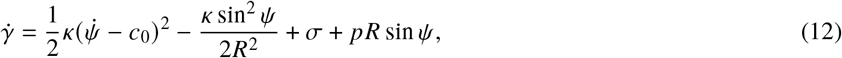

and

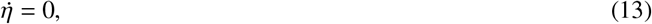

as well as Eqs. (7) and (8).

For the homogeneous model, the spontaneous curvature *c*_0_ is uniform and 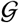 is explicitly independent of the arclength *s*. This symmetry leads to a conserved quantity (55)

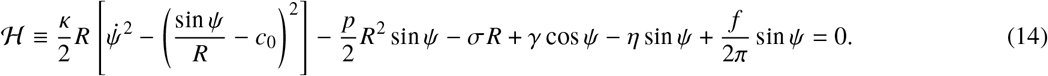

For the inhomogeneous model, the spontaneous curvature *c*_0_ [*a*(*s*)] is spatially varied over the arclength *s* as depicted by Eq. (6). The variation of the functional *G* in Eq. (10) needs to change to include a spatially varied surface tension *σ*(*S*) to ensure that the membrane area is locally unstretchable. The detailed derivation can be found in Ref. (16). The equation for *σ* reads

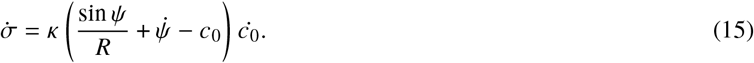

In addition to the varied surface tension, Eq. (11) for the membrane angle *ψ* needs to change to include a new term 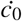,

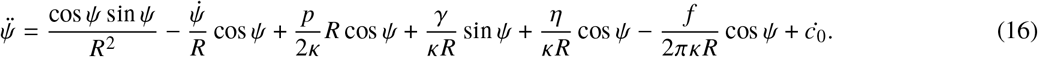

It is easy to verify that the new Eqs.(15) and (16) together with Eqs. (12) and (13) ensure that 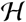 is conserved, i.e., 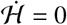. When considering the effect of distributed forces, we assume the forces are localized in an area of *a*_f_ near the membrane tip and pointing in the normal direction. Specifically, we express the normal stress (force per unit area) *g_n_* as the following form

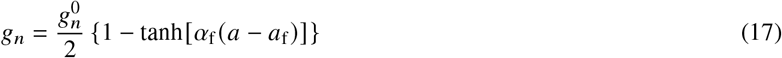

where 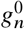 denotes the magnitude of the normal stress within the area *a*_f_ where forces are loaded, and *α*_f_ controls the sharpness of the force drop at *a*_f_. In this case, Eqs. (12) and (13) need to make the following changes,

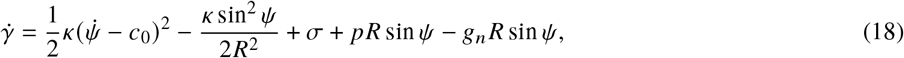

and

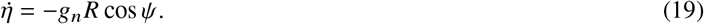

The total force *f* is calculated as the surface integral of the normal stress

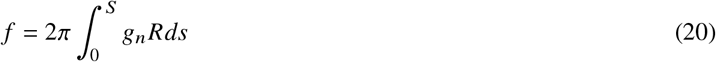

### Derivation of the boundary conditions

In order to get proper BCs, we set the boundary terms in Eq. (10) to zero. At the membrane tip (*s* = 0), *R* = 0 by definition and we choose *ψ* = 0 to avoid any singularity. As a result, *δR* = 0 and *δψ* = 0 and the boundary terms automatically vanish.

At the base of the invagination (*s* = *S*), as a result of the product of two conjugate variables and 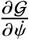 and *δψ*, we have the freedom to let either 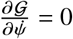, i.e. the membrane can be freely rotate (free-hinge BC), or *δψ =* 0, i.e. the angle of the membrane is fixed (fixed-hinge BC). Similarly, we can choose 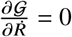, i.e. the base can freely move, or *δR* = 0, i.e. the base radius is fixed. The combination of the two choices make up the four possible BCs listed in Table 1.

### Numerical methods to calculate the force-height (*f-L*) relationships

For the homogeneous model with a uniform spontaneous curvature *c*_0_, Eqs. (7), (11), (12) and *η* = 0 constitute a complete system of equations, which are numerically solved by a shooting method that has been widely used in Helfrich models (18, 50). The idea is to numerically integrate the three equations from the membrane tip *s* = 0 with MATLAB solver ode45 until the free-hinge BC or the fixed-hinge BC is met. The numerical integration needs input of the initial values of *R*(*s* = 0), *ψ*(*s* = 0), 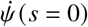 and *γ*(*s* = 0). The radius *R*(*s* = 0) should be zero at the membrane tip. However, Eqs. (11) and (12) have a singular point at *R* = 0. In order to avoid the singular point, we set *R*(*s* = 0) = *ϵR*_p_, where *ϵ* = 0.001 is chosen to be a small number such that values smaller than 0.001 do not produce numerically distinguishable results. The initial angle *ψ*(*s* = 0) = 0 is to ensure continuity of the membrane shape at the tip. The derivative 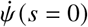 is the tuning parameter to match the BCs. For any given 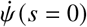, *γ*(*s* = 0) is solved via Eq. (14). Once the four initial values are set, the numerical integration continues until the free-hinge BC or the fixed-hinge BC is met. This is achieved by setting the termination event function in the ode45 solver. The membrane height 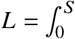 is then obtained via Eq. (8). Note that for different trials, the final arclength *S* when the solver terminates are different. The shooting method is to find a proper pair of 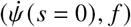 such that when the integration terminates, i.e., the free-hinge BC or the fixed-hinge has been satisfied, the other BCs *R* = *R*_b_ and *L* = *L*_0_ are fulfilled for a particular membrane height *L*_0_. In order to construct the *f-L* curve, once we get the solution of 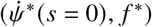 for a particular *L*_0_, we extend the membrane height *L* with a small increment to *L*_0_ + Δ*L*. The solution 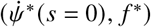 for *L* = *L*_0_ are then used as the initial trial for searching the solution for *L* = *L*_0_ + Δ*L*.

For the inhomogeneous model with a spatially varied spontaneous curvature *c*_0_ (*S*) defined by Eq. (6), Eqs. (7), (12), (15), (16) and 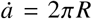, *η* = 0 constitute a complete system of equations. In addition to the four initial values required by the homogeneous model, *a*(*s* = 0) and *σ*(*s* = 0) are needed to numerically integrate the equations. We set *a*(*s* = 0) = 0 and tune the combination of 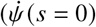, *σ*(*s* = 0), *f*) to match *R* = *R*_b_, *L* = *L*_0_ and *σ* = *σ*_0_ when the solver terminates. The *f-L* curve is constructed in a similar way by gradually extending the membrane height L with small increment of Δ*L*.

When considering the effect of distributed forces on a homogeneous membrane, Eqs. (7), (11) (with *f* = 0), (18), (19), and 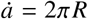 constitute a complete system of equations. In addition to the four initial values required by the homogeneous model, *a*(*s* = 0) and *η*(*s* = 0) are needed to numerically integrate the equations. We set *a*(*s* = 0) = 0, *η*(*s* = 0) = 0 and tune the combination of 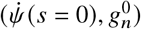 to match *R* = *R*_b_, *L* = *L*_0_ when the solver terminates. The total force is calculated with (20).

### Numerical procedure to fit the experimental data

We have 7 parameters in the inhomogeneous model listed in Table 2. The turgor pressure *p* is fixed at *p* = 1MPa. For the remaining 6 parameters, we express five of them as the function of the characteristic radius *R*_p_ and use *R*_p_ as the single parameter to fit the experimental data. The surface tension at the base *σ* is set to be 0.002*pR*_p_ such that the surface tension σ plays a much less important role than the turgor pressure *p* in determining the tube radius because 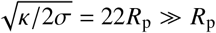. The base radius *R*_b_ is fixed at *R*_b_ = 2*R*_p_ such that for a bare membrane, the force barrier *F*_max_ as a function of *R*_b_ is close to the plateau and not sensitive to the variation of base radius (see Figure2 e and f). Based on the experimental observation that the copy number of clathrin molecules stays small and almost constant during the assembly and disassembly of actin meshwork (46), and the measured copy number 30-40 implies a hemispherical cap of the clathrin coat, we assume the coating area 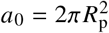 and the spontaneous curvature *c*_0_ = 1/*R*_p_. The sharpness of the coating edge is controlled by the parameter *α*, which is set to be 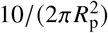. Values of *α* greater than 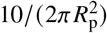 do not make a difference on the resulting *f-L* curve (Figure S7).

We use the geometric features *R*_t_ and *D*_t_ v.s. membrane height L as our fitting data. For the data points of 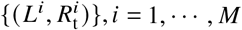 in Figure 6c and d, the corresponding theoretical prediction of the tip radius ThR(*L^i^*) is calculated for a given *R*_p_. The fitting error then reads

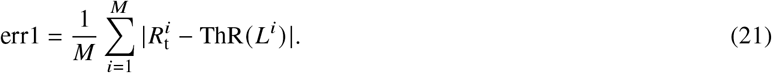

Similarly the fitting error for the distance from neck to tip *D_t_* reads

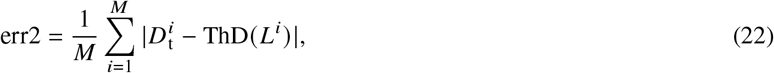

where ThD(*L^i^*) denotes the theoretical prediction of *D*_t_ at *L* = *L^i^*. When plotting err1 + err2 as a function of the fitting parameter *R*_p_, we find the optimum 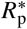 that minimizes the sum err1 + err2 (Figure S4).

## AUTHOR CONTRIBUTIONS

RM and JB designed the research. RM carried out all simulations, analyzed the data. RM and JB wrote the article.

## ACKNOWLEDGMENTS

This research is supported by National Institutes of Health/National Institute of General Medical Sciences Grant R01GM115636. We thank Wanda Kukulski, Marko Kaksonen and John Briggs for kindly sharing the micrograph of Figure 1a. We thank Dr. Pablo Sartori for critical reading of the manuscript.

## SUPPLEMENTARY MATERIAL

An online supplement to this article can be found by visiting BJ Online at http://www.biophysj.org.

## Supporting Material for

### DERIVATION OF THE ANALYTICAL SOLUTION FOR THE INITIATION FORCE

The derivation here is based on the analytical solution obtained in the work of (1) in the limit of small angles. It assumes that the membrane is almost flat such that the tangential *ψ* ≪ 1. Keeping only the first order of *ψ* and its derivatives and performing the coordinate transformation of *ψ* (*s*) to *ψ*(*R*), the shape equation can be reduced to

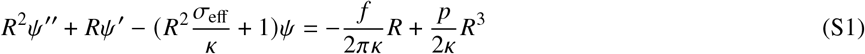

where the prime (′) indicates the derivative with respect to the radial coordinate *R* and 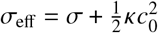.

The general solution to Eq. (S1) that satisfies the boundary condition at the tip *ψ*(*R* = 0) = 0 reads

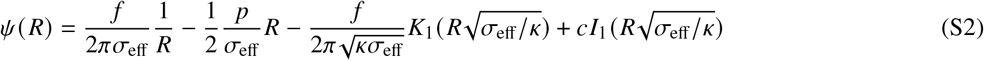

where *I_i_*(*x*) and *K_i_*(*x*) are modified Bessel functions. The constant c is determined by the other boundary condition at *R* = *R*_b_. For the free-hinge BC, it reads

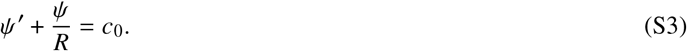

For fixed-hinge BC, it reads

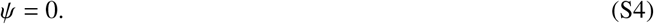

To calculate the initiation force *F*_init_, we set the membrane height *L* = 0, i.e.,

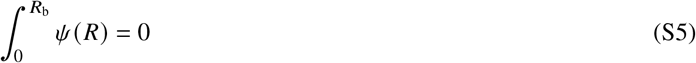

For the free-hinge BC, the initiation force in the limit of *σ* → 0 reads

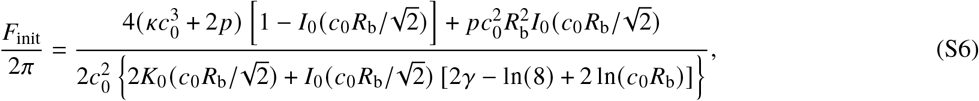

where *γ* is the Euler constant. As a special case *c*_0_ → 0, Eq. (S6) reduces to

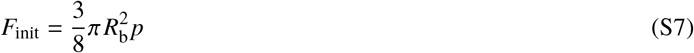

For the fixed-hinge BC, the initiation force reads

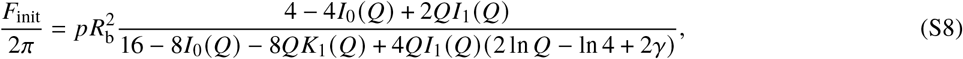

where 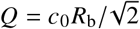. As a special case *c*_0_ → 0, Eq. (S8) reduces to

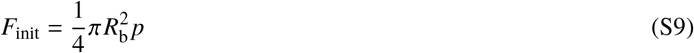

### SPHERICAL CAP APPROXIMATION

We approximate the dome-shaped membrane with a spherical cap of radius *R* (see Figure 7, a and b, inset). The shape of the spherical cap is fully determined by two parameters, the sphere radius *R* and the base radius *R*_b_. Note that neither the free-hinge BC nor the fixed-hinge BC is satisfied in this case. The membrane height *L* of the spherical cap reads

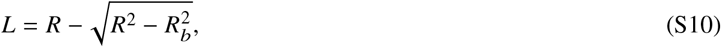

and the corresponding membrane area *A* = 2*πRL* and the volume 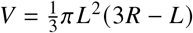. The free energy of the membrane then becomes

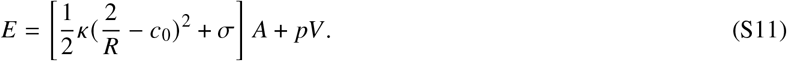

In the case of a fixed base radius, the free energy *E*(*R*; *c*_0_, *R*_b_) is a function of only *R*. For a barely coated membrane, i.e., *c*_0_*R*_p_ = 0, the free energy *E*(*R*; *c*_0_, *R*_b_) decreases monotonically with the radius *R* and the flat shape (*R* = ∞) has the lowest energy (Figure 7 a). However, when *c*_0_ becomes large, e.g., *c*_0_*R*_p_ = 1, the free energy *E*(*R*; *c*_0_, *R*_b_) has a minimum at a finite radius *R* (Figure 7 b). This means the flat shape is no longer stable and the membrane can be bent up by proteins without any external forces.

If the base radius is free to move, i.e., *R*_b_ being a free parameter, the free energy *E*(*R*, *R*_b_; *c*_0_) becomes a function of both *R* and *R*_b_, and always has its trivial minimum *E* = 0 at *R*_b_ = 0, regardless of the spontaneous curvature *c*_0_ (Figure 7a and b, *R*_b_ = 0*R*_p_). The corresponding solution at *R*_b_ = 0 represents an infinitely small patch of membrane. In the presence of an external force *f*, the total free energy *F*(*R*, *R*_b_; *c*_0_, *f*) = *E*(*R*, *R*_b_; *c*_0_, *f*) - *fL* can have a nontrivial minimum with a nonzero *R*_b_. The minimum force *f*_min_ to have such a nontrivial minimum, i.e., to lift the membrane up, is given by

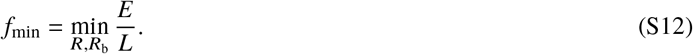

Both the denominator and the numerator are positive numbers, therefore *f*_min_ is always positive. For instance, for *c*_0_*R*_p_ = 1, the minimum force *f*_min_ = 0.0027*f*_p_ is obtained at *R* = 1.998*R*_p_ and *R*_b_ = 0.00057*R*_p_. Therefore an external force is always required to lift the membrane up in this condition. However, the base radius *R*_b_ = 0.00057*R*_p_ is an unrealistically narrow shape given the typical value of *R*_p_ = 15-30 nm. Similar problem also exists in the model of (2) which used a freely-moving base BC.

## SUPPLEMENTAL FIGURES

**Figure S1:**
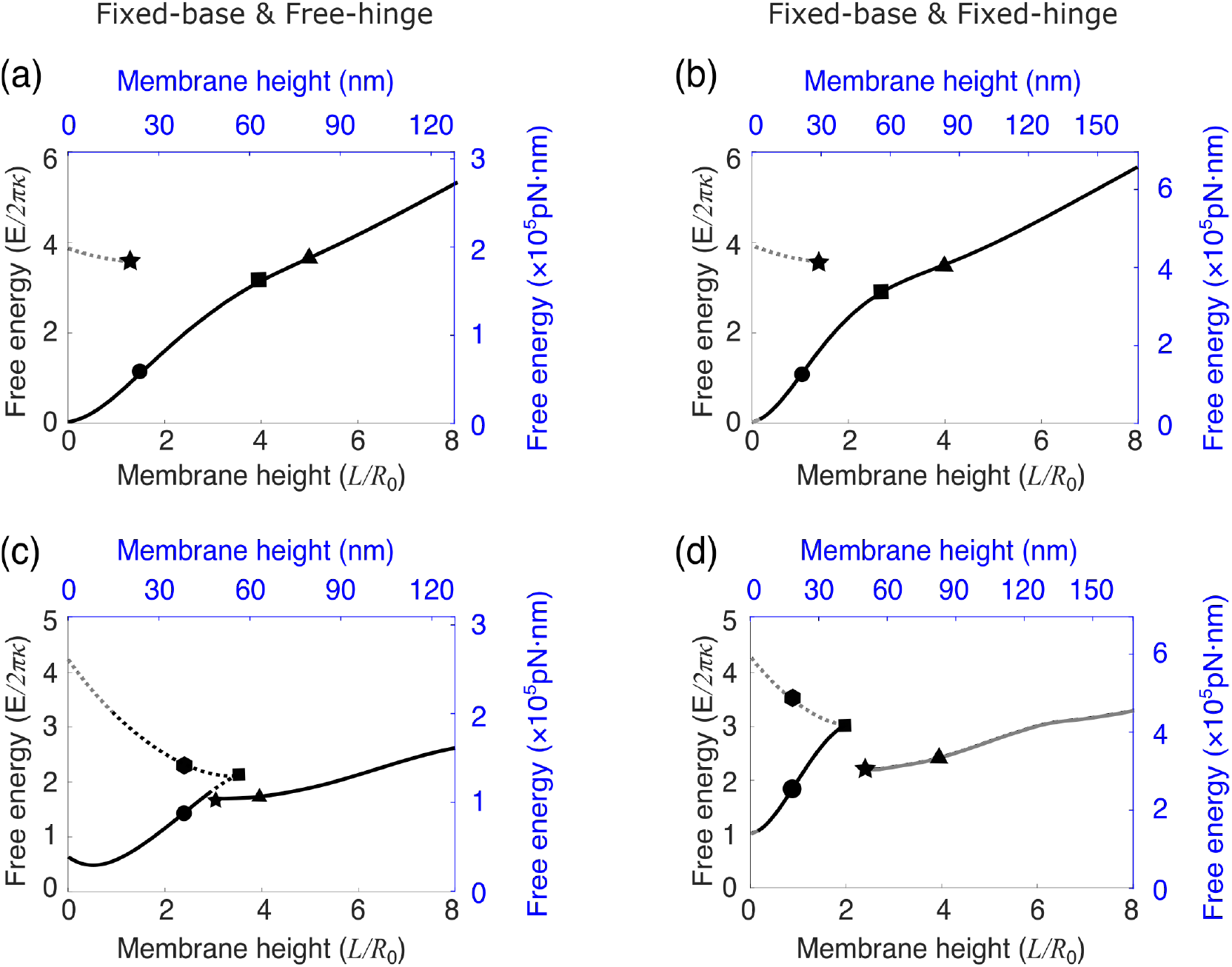
Free energy of membrane deformations for a fully coated membrane. (a - d) Free energy of membrane deformations *E* as a function of membrane height L. The spontaneous curvature is fixed at *c*_0_*R*_p_ = 0.2 in (a, b) and *c*_0_*R*_p_ = 1 in (c, d). The symbols on the E-L curve correspond to the same symbols on the *f-L* curve shown in Figure 3a-d. In (a - d), the solid line indicates shapes of the lowest free energy and the dashed line indicates shapes of relatively high free energy. The dark color indicates membrane shapes that are all above *z =* 0, and the gray color indicates shapes that have parts below *z* = 0. In the left column (a, c), the free-hinge BC is imposed at the base points *R*_b_ = 2*R*_p_, while in the right column (b, d), the fixed-hinge BC is imposed. On the left and bottom axes (black), non-dimensionalized quantities are used, while on the right and top axes (blue), quantities are measured in their physical units. The parameters are listed in TABLE 2.

**Figure S2:**
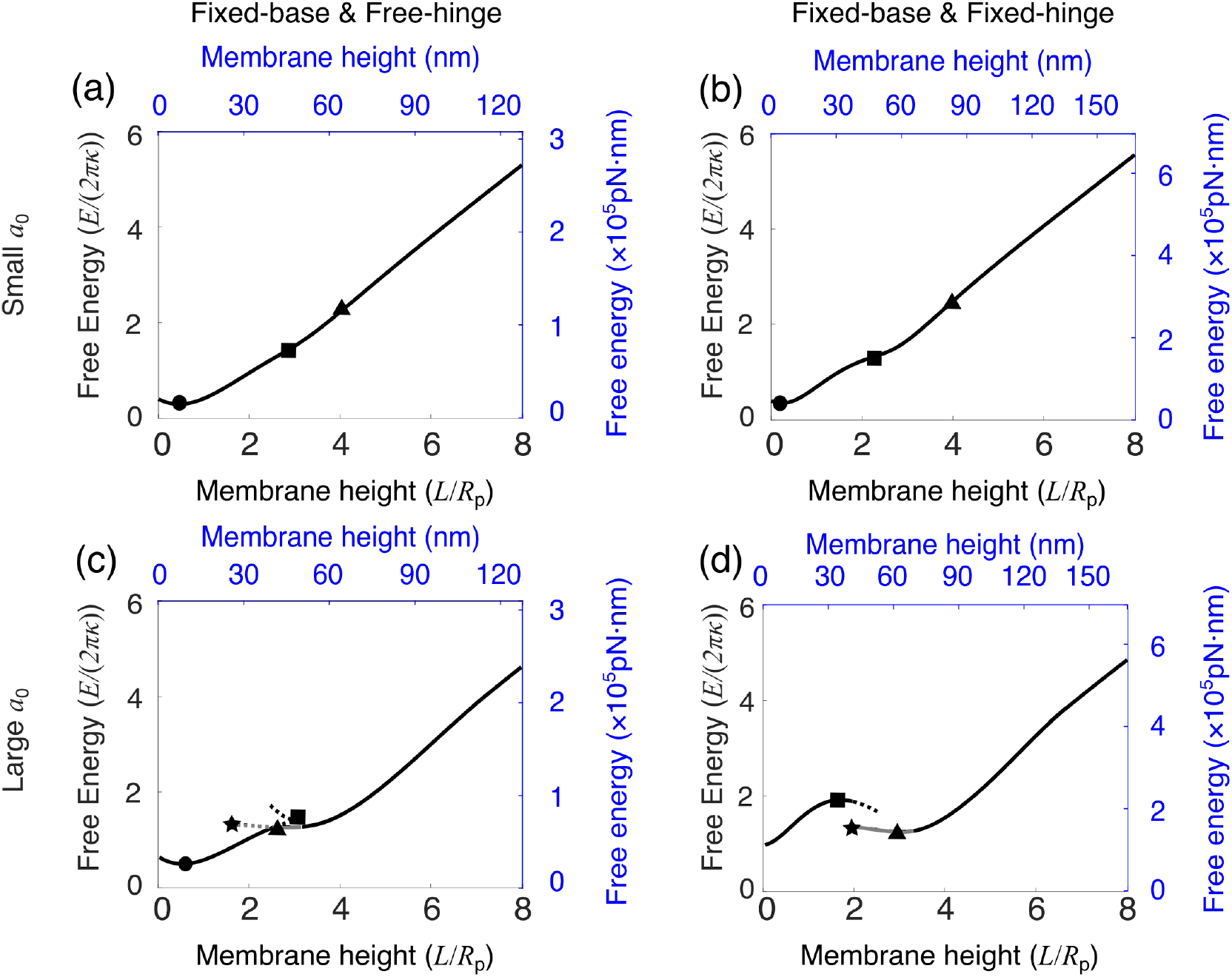
Free energy of membrane deformations for a partially coated membrane. (a - d) Free energy of membrane deformations E as a function of membrane height L. The coating area is fixed at 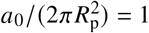 in (a, b) and 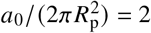 in (c, d). The symbols on the E-L curve correspond to the same symbols on the *f-L* curve shown in Figure 4a-d. In (a - d), the solid line indicates shapes of the lowest free energy and the dashed line indicates shapes of relatively high free energy. The dark color indicates membrane shapes that are all above *z* = 0, and the gray color indicates shapes that have parts below *z* = 0. In the left column (a, c), the free-hinge BC is imposed at the base points *R*_b_ = 2*R*_p_, while in the right column (b, d), the fixed-hinge BC is imposed. On the left and bottom axes (black), non-dimensionalized quantities are used, while on the right and top axes (blue), quantities are measured in their physical units. The parameters are listed in TABLE 2.

**Figure S3:**
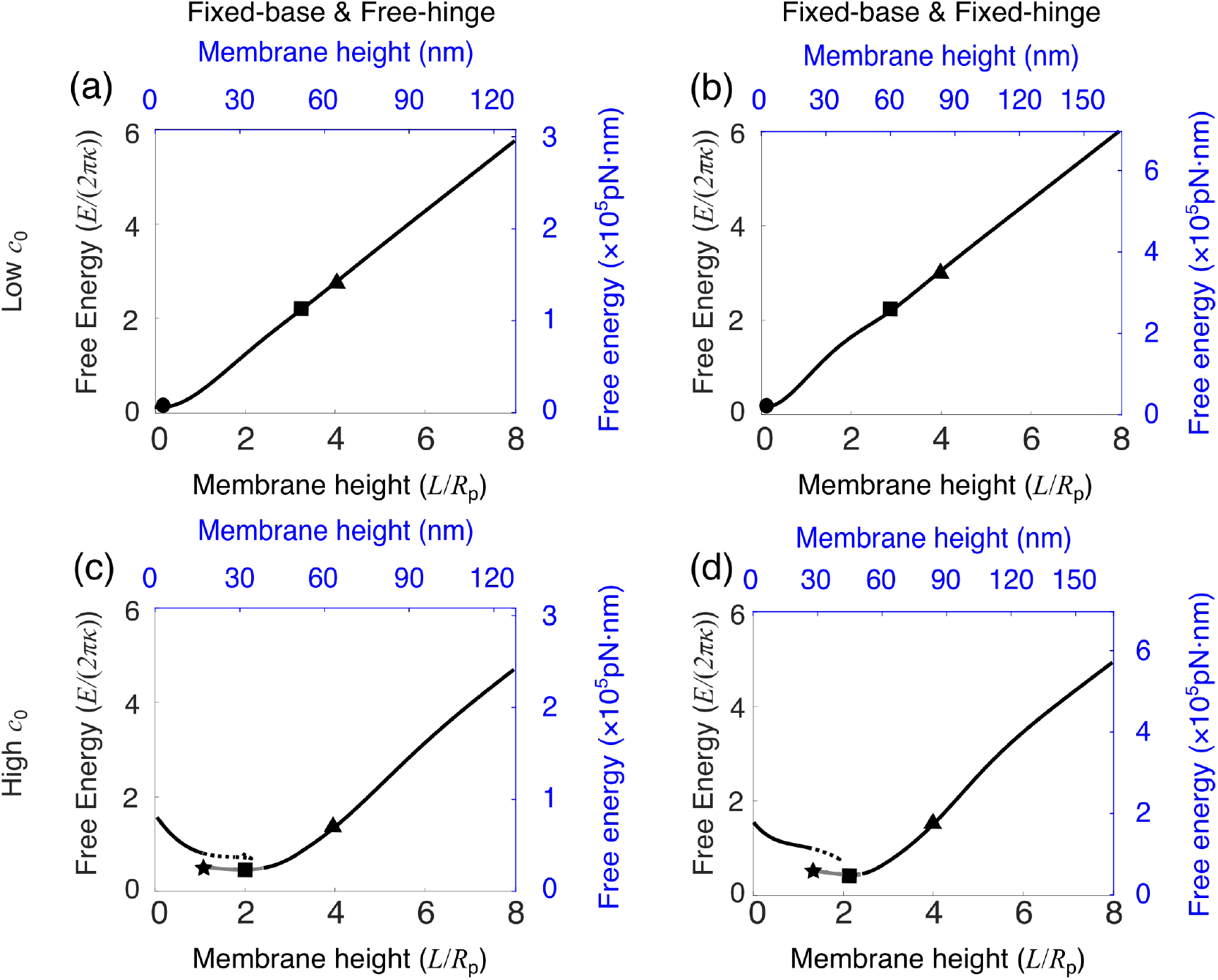
Free energy of membrane deformations for a partially coated membrane. (a - d) Free energy of membrane deformations E as a function of membrane height L. The coating area is fixed at 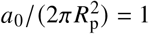 in (a, b) and 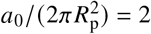 in (c, d). The symbols on the E-L curve correspond to the same symbols on the *f-L* curve shown in Figure 5a-d. In (a - d), the solid line indicates shapes of the lowest free energy and the dashed line indicates shapes of relatively high free energy. The dark color indicates membrane shapes that are all above *z* = 0, and the gray color indicates shapes that have parts below *z* = 0. In the left column (a, c), the free-hinge BC is imposed at the base points *R*_b_ = 2*R*_p_, while in the right column (b, d), the fixed-hinge BC is imposed. On the left and bottom axes (black), non-dimensionalized quantities are used, while on the right and top axes (blue), quantities are measured in their physical units. The parameters are listed in TABLE 2.

**Figure S4:**
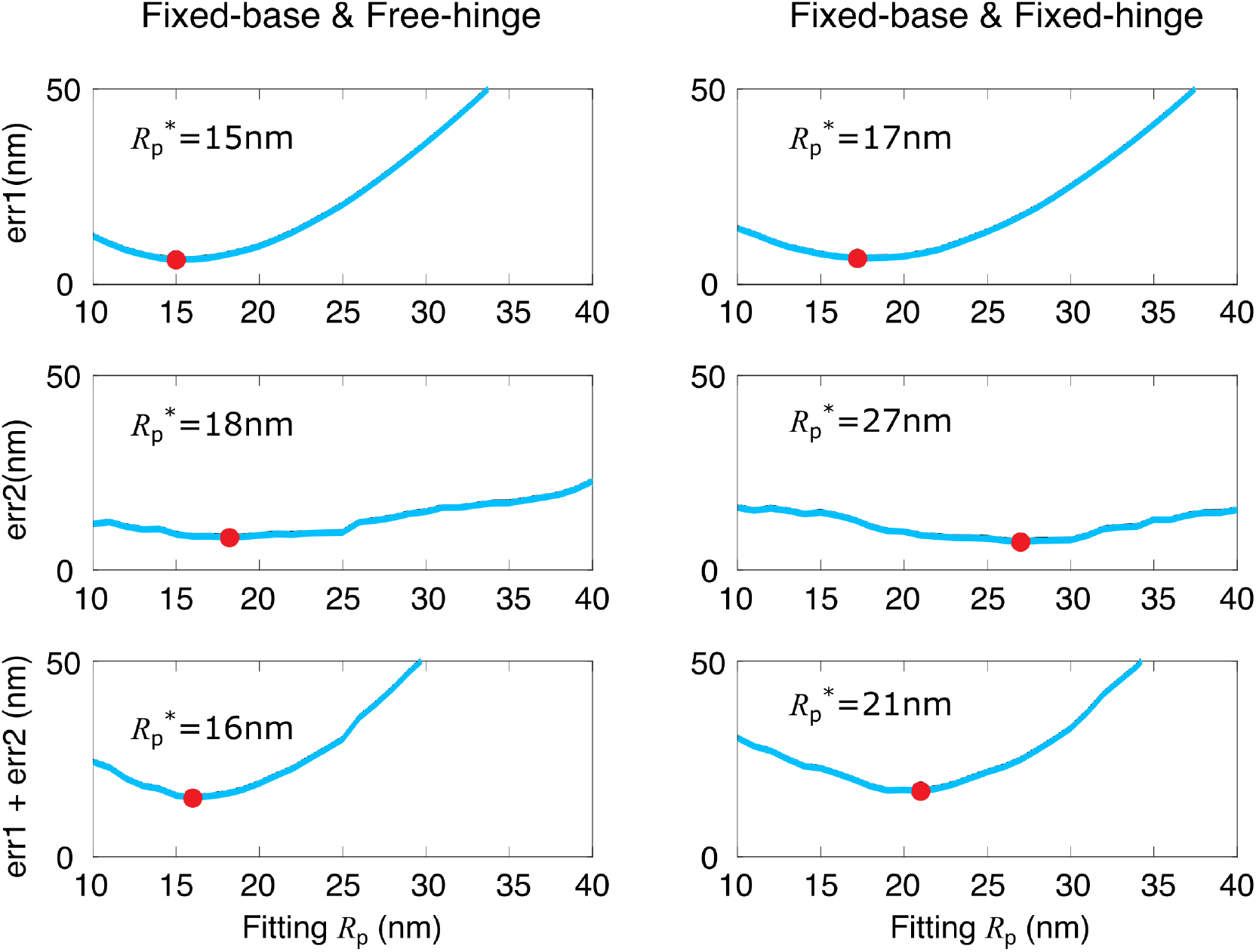
The fitting error as a function of the fitting parameter *R*_p_ for the two BCs. The first row shows the fitting error for the dataset of *R*_t_ v.s. L, and the second row for the dataset of *D*_t_ v.s. L and the third row for the sum. In the left column, the free-hinge BC is imposed, while in the right column, the fixed-hinge BC is imposed.

**Figure S5:**
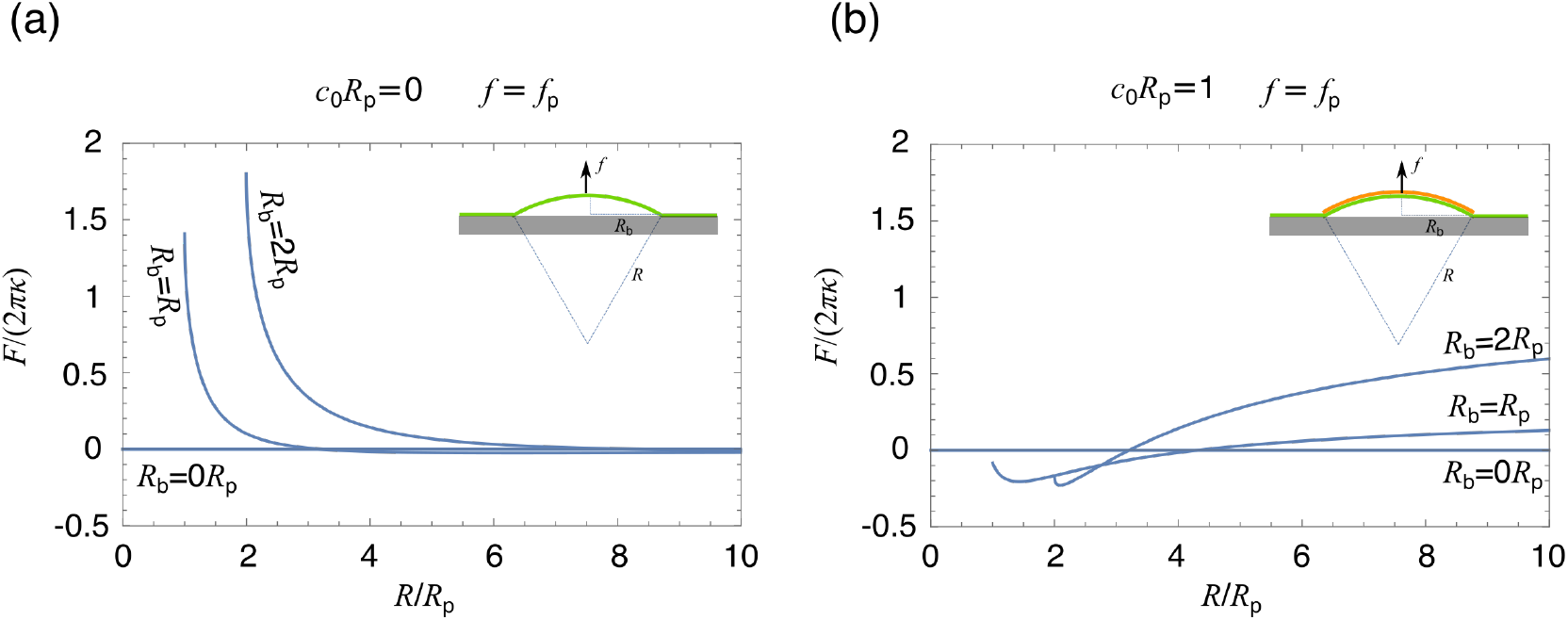
Free energy of membrane deformations in the presence of external force under spherical cap approximation. Free energy of the membrane as a function of the sphere radius R for *c*_0_*R*_p_ = 0 in (a) and *c*_0_*R*_p_ = 1 in (b). The external force *f =* f_p_. For different base radii *R*_b_, the range of R is [*R*_b_, ∞], where *R* = *R*_b_ corresponds to a hemi-spherical cap and *R* = ∞ corresponds to a flat shape.

**Figure S6:**
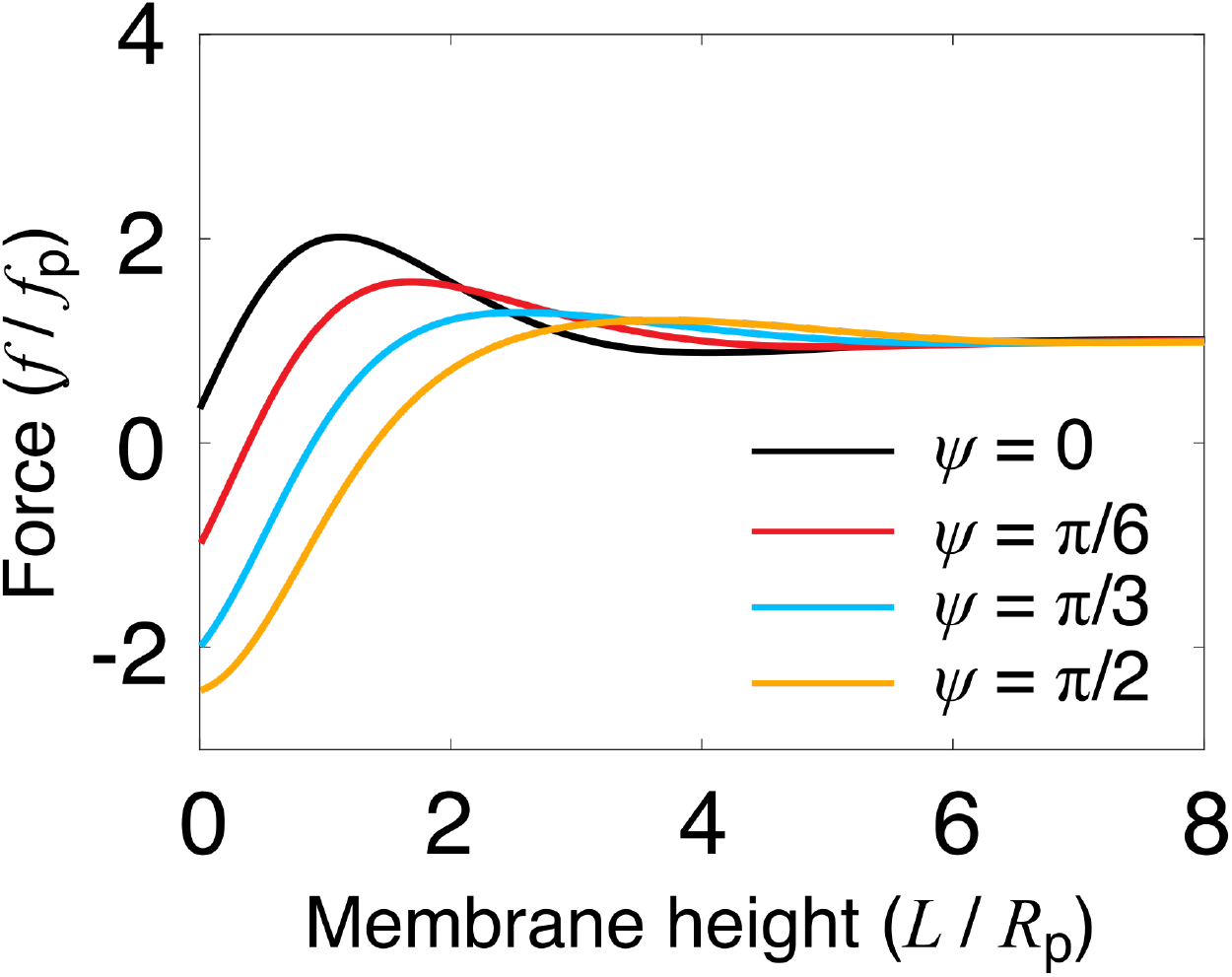
*F-L* curves for the fixed-hinge BC with different values of the angle *ψ* at the base for a fully uncoated membrane. The parameters are the same as in Figure 2d except that the angle at the base is varied.

**Figure S7:**
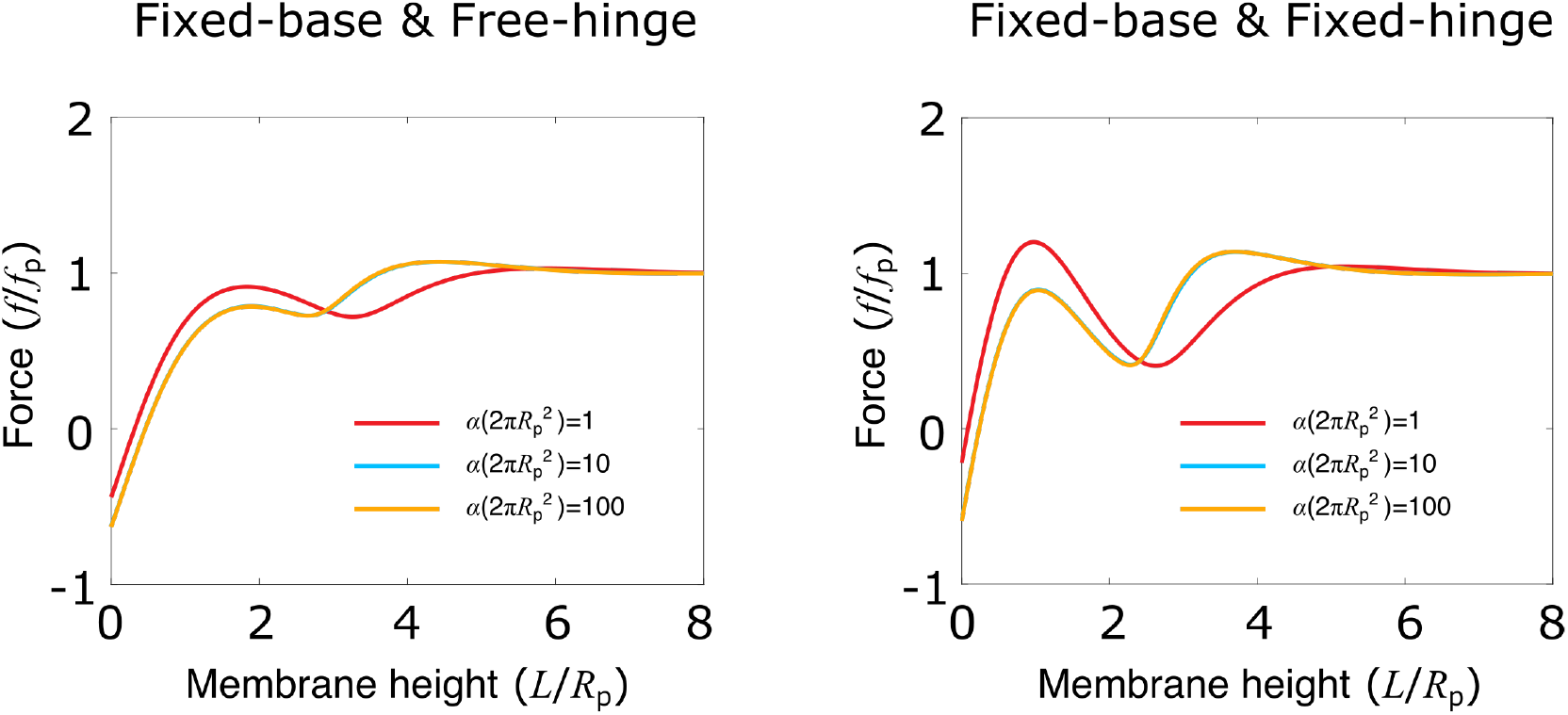
*F-L* curves for a partially coated membrane with different values of *α* that controls the sharpness of the coating edge. The parameters are the same as in Figure 4a and b except that the parameter *α* that controls the sharpness of the coating edge is varied. In the left column, the free-hinge BC is imposed at the base points *R*_b_ = 2*R*_p_, while in the right column, the fixed-hinge BC is imposed.

**Figure S8:**
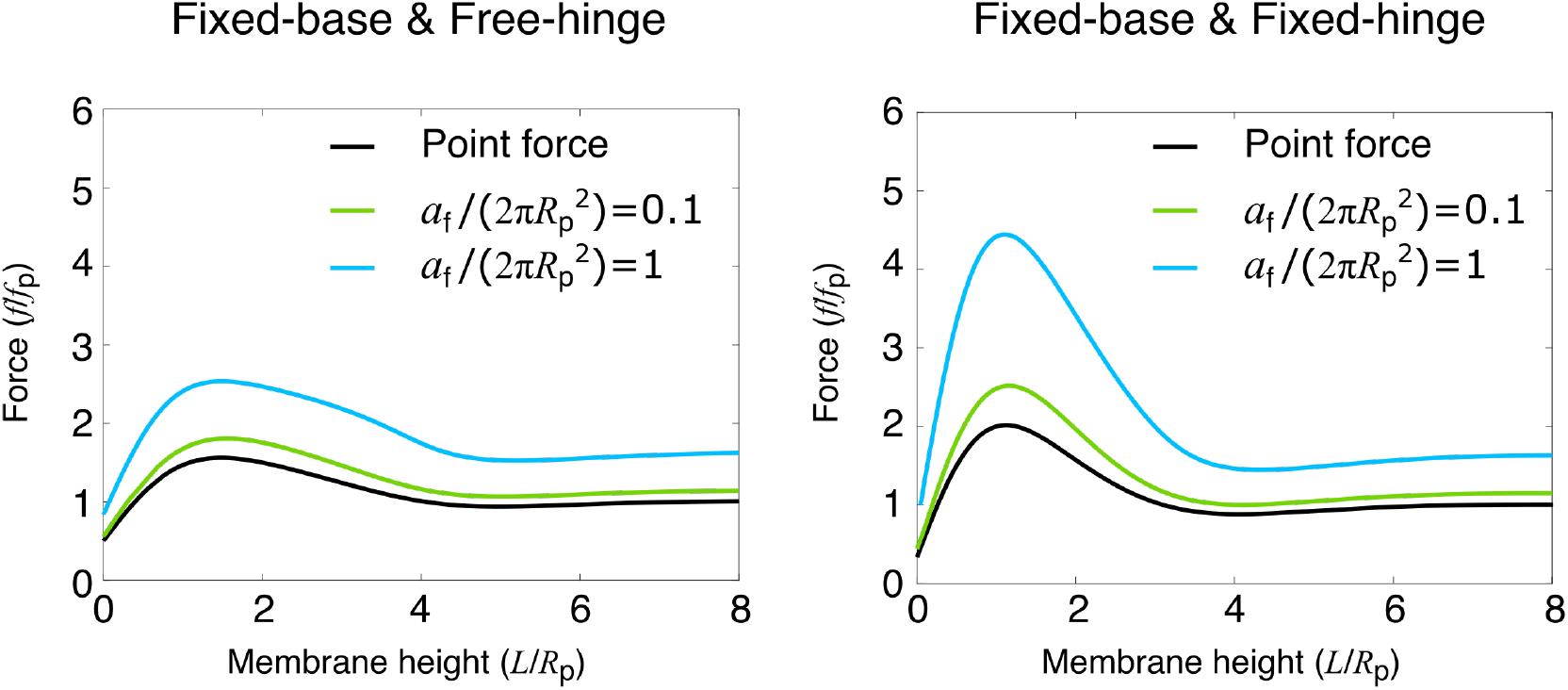
*F-L* curves for distributed forces for a bare membrane. The forces are assumed to be distributed in an area of *a_f_* near the membrane tip and pointing in the normal direction. The black curve represents the *f-L* curve under the point force assumption. The parameters are the same as in Figure 2c and d with an additional parameter 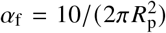. In the left column, the free-hinge BC is imposed at the base points *R*_b_ = 2*R*_p_, while in the right column, the fixed-hinge BC is imposed.

## Notes

https://github.com/ruima86/MembraneModel

### Competing Interest Statement

The authors have declared no competing interest.

## REFERENCES

1. McMahon, H. T., and E. Boucrot, 2011. Molecular mechanism and physiological functions of clathrin-mediated endocytosis. Nature Reviews Molecular Cell Biology 12:517–533. https://doi.org/10.1038/nrm3151.

2. Sorkin, A., and M. A. Puthenveedu, 2013. Clathrin-Mediated Endocytosis, Springer New York, New York, NY, 1–31. https://doi.org/10.1007/978-1-4614-6528-7_1.

3. Lu, R., D. G. Drubin, and Y. Sun, 2016. Clathrin-mediated endocytosis in budding yeast at a glance. Journal of Cell Science 129:1531–1536. https://jcs.biologists.org/content/129/8/1531.

4. Kaksonen, M., and A. Roux, 2018. Mechanisms of clathrin-mediated endocytosis. Nature Reviews Molecular Cell Biology 19:313 EP –. https://doi.org/10.1038/nrm.2017.132.

5. Lacy, M. M., R. Ma, N. G. Ravindra, and J. Berro, 2018. Molecular mechanisms of force production in clathrin-mediated endocytosis. FEBS Letters 0. https://febs.onlinelibrary.wiley.com/doi/abs/10.1002/1873-3468.13192.

6. Mettlen, M., P.-H. Chen, S. Srinivasan, G. Danuser, and S. L. Schmid, 2018. Regulation of Clathrin-Mediated Endocytosis. Annual Review of Biochemistry 87:871–896. https://doi.org/10.1146/annurev-biochem-062917-012644, pMID: 29661000.

7. Boulant, S., C. Kural, J.-C. Zeeh, F. Ubelmann, and T. Kirchhausen, 2011. Actin dynamics counteract membrane tension during clathrin-mediated endocytosis. Nature Cell Biology 13:1124–1131. https://doi.org/10.1038/ncb2307.

8. Wu, X.-S., S. Elias, H. Liu, J. Heureaux, P. J. Wen, A. P. Liu, M. M. Kozlov, and L.-G. Wu, 2017. Membrane Tension Inhibits Rapid and Slow Endocytosis in Secretory Cells. Biophysical Journal 113:2406–2414. http://www.sciencedirect.com/science/article/pii/S0006349517310810.

9. Low, P. S., and S. Chandra, 1994. Endocytosis in plants. Annual review of plant biology 45:609–631.

10. Aghamohammadzadeh, S., and K. R. Ayscough, 2009. Differential requirements for actin during yeast and mammalian endocytosis. Nature Cell Biology 11:1039–1042. https://doi.org/10.1038/ncb1918.

11. Basu, R., E. L. Munteanu, and F. Chang, 2014. Role of turgor pressure in endocytosis in fission yeast. Molecular biology of the cell 25:679–687. https://www.ncbi.nlm.nih.gov/pubmed/24403609.

12. Minc, N., A. Boudaoud, and F. Chang, 2009. Mechanical Forces of Fission Yeast Growth. Current Biology 19:1096–1101. http://www.sciencedirect.com/science/article/pii/S0960982209011324.

13. Atilgan, E., V. Magidson, A. Khodjakov, and F. Chang, 2015. Morphogenesis of the Fission Yeast Cell through Cell Wall Expansion. Current Biology 25:2150–2157. https://doi.org/10.1016/j.cub.2015.06.059.

14. Kukulski, W., M. Schorb, M. Kaksonen, and J. A. Briggs, 2012. Plasma Membrane Reshaping during Endocytosis Is Revealed by Time-Resolved Electron Tomography. Cell 150:508–520. http://www.sciencedirect.com/science/article/pii/S0092867412007842.

15. Avinoam, O., M. Schorb, C. J. Beese, J. A. G. Briggs, and M. Kaksonen, 2015. Endocytic sites mature by continuous bending and remodeling of the clathrin coat. Science 348:1369–1372. http://science.sciencemag.org/content/348/6241/1369.

16. Agrawal, A., and D. J. Steigmann, 2008. Modeling protein-mediated morphology in biomembranes. Biomechanics and Modeling in Mechanobiology 8:371. https://doi.org/10.1007/s10237-008-0143-0.

17. Walani, N., J. Torres, and A. Agrawal, 2015. Endocytic proteins drive vesicle growth via instability in high membrane tension environment. Proceedings of the National Academy of Sciences 112:E1423–E1432. http://www.pnas.org/content/112/12/E1423.

18. Dmitrieff, S., and F. Nédélec, 2015. Membrane Mechanics of Endocytosis in Cells with Turgor. PLoS Comput Biol 11:1–15. http://dx.doi.org/10.1371%2Fjournal.pcbi.1004538.

19. Hassinger, J. E., G. Oster, D. G. Drubin, and P. Rangamani, 2017. Design principles for robust vesiculation in clathrin- mediated endocytosis. Proceedings of the National Academy of Sciences 114: E1118–E1127. http://www.pnas.org/content/114/7/E1118.

20. Alimohamadi, H., R. Vasan, J. E. Hassinger, J. C. Stachowiak, and P. Rangamani, 2018. The role of traction in membrane curvature generation. Molecular biology of the cell 29:2024–2035.

21. Napoli, G., and A. Goriely, 2020. Elastocytosis. Journal of the Mechanics and Physics of Solids 145:104133.

22. Koster, G., A. Cacciuto, I. Derényi, D. Frenkel, and M. Dogterom, 2005. Force Barriers for Membrane Tube Formation. Phys. Rev. Lett. 94:068101. https://link.aps.org/doi/10.1103/PhysRevLett.94.068101.

23. Cuvelier, D., I. Derényi, P. Bassereau, and P. Nassoy, 2005. Coalescence of Membrane Tethers: Experiments, Theory, and Applications. Biophysical Journal 88:2714–2726. http://www.sciencedirect.com/science/article/pii/S0006349505733257.

24. Dimova, R., S. Aranda, N. Bezlyepkina, V. Nikolov, K. A. Riske, and R. Lipowsky, 2006. A practical guide to giant vesicles. Probing the membrane nanoregime via optical microscopy. Journal of Physics: Condensed Matter 18:S1151. http://stacks.iop.org/0953-8984/18/i=28/a=S04.

25. Zhong-Can, O.-Y., and W. Helfrich, 1987. Instability and deformation of a spherical vesicle by pressure. Physical review letters 59:2486.

26. Seifert, U., K. Berndl, and R. Lipowsky, 1991. Shape transformations of vesicles: Phase diagram for spontaneous-curvature and bilayer-coupling models. Physical review A 44:1182.

27. Seifert, U., 1997. Configurations of fluid membranes and vesicles. Advances in physics 46:13–137.

28. Mund, M., J. A. van der Beek, J. Deschamps, S. Dmitrieff, P. Hoess, J. L. Monster, A. Picco, F. Nédélec, M. Kaksonen, and J. Ries, 2018. Systematic Nanoscale Analysis of Endocytosis Links Efficient Vesicle Formation to Patterned Actin Nucleation. Cell 174:884–896.e17. http://www.sciencedirect.com/science/article/pii/S0092867418308006.

29. Carlsson, A. E., 2018. Membrane bending by actin polymerization. Current Opinion in Cell Biology 50:1–7. http://www.sciencedirect.com/science/article/pii/S095506741730128X, cell Architecture.

30. Kübler, E., and H. Riezman, 1993. Actin and fimbrin are required for the internalization step of endocytosis in yeast. The EMBO journal 12:2855–2862.

31. Engqvist-Goldstein, Å. E., and D. G. Drubin, 2003. Actin Assembly and Endocytosis: From Yeast to Mammals. Annual Review of Cell and Developmental Biology 19:287–332. https://doi.org/10.1146/annurev.cellbio.19.111401.093127, pMID: 14570572.

32. Yarar, D., C. M. Waterman-Storer, and S. L. Schmid, 2005. A Dynamic Actin Cytoskeleton Functions at Multiple Stages of Clathrin-mediated Endocytosis. Molecular Biology of the Cell 16:964–975. https://doi.org/10.1091/mbc.e04-09-0774, pMID: 15601897.

33. Sun, Y., A. C. Martin, and D. G. Drubin, 2006. Endocytic Internalization in Budding Yeast Requires Coordinated Actin Nucleation and Myosin Motor Activity. Developmental Cell 11:33–46. http://www.sciencedirect.com/science/article/pii/S1534580706002462.

34. Kaksonen, M., C. P. Toret, and D. G. Drubin, 2006. Harnessing actin dynamics for clathrin-mediated endocytosis. Nature Reviews Molecular Cell Biology 7:404–414. https://doi.org/10.1038/nrm1940.

35. Mooren, O. L., B. J. Galletta, and J. A. Cooper, 2012. Roles for actin assembly in endocytosis. Annual review of biochemistry 81:661–686.

36. Goode, B. L., J. A. Eskin, and B. Wendland, 2015. Actin and endocytosis in budding yeast. Genetics 199:315–358.

37. Berro, J., and T. D. Pollard, 2014. Local and global analysis of endocytic patch dynamics in fission yeast using a new “temporal superresolution” realignment method. Molecular Biology of the Cell 25:3501–3514. https://doi.org/10.1091/mbc.e13-01-0004, pMID: 25143395.

38. Carlsson, A. E., and P. V. Bayly, 2014. Force Generation by Endocytic Actin Patches in Budding Yeast. Biophysical Journal 106:1596–1606. http://www.sciencedirect.com/science/article/pii/S0006349514002823.

39. Wang, X., B. J. Galletta, J. A. Cooper, and A. E. Carlsson, 2016. Actin-Regulator Feedback Interactions during Endocytosis. Biophysical Journal 110:1430–1443. http://www.sciencedirect.com/science/article/pii/80006349516001648.

40. Tweten, D. J., P. V. Bayly, and A. E. Carlsson, 2017. Actin growth profile in clathrin-mediated endocytosis. Phys. Rev. E 95:052414. https://link.aps.org/doi/10.1103/PhysRevE.95.052414.

41. Kirchhausen, T., and S. C. Harrison, 1981. Protein organization in clathrin trimers. Cell 23:755–761. http://www.sciencedirect.com/science/article/pii/0092867481904396.

42. Fotin, A., Y. Cheng, P. Sliz, N. Grigorieff, S. C. Harrison, T. Kirchhausen, and T. Walz, 2004. Molecular model for a complete clathrin lattice from electron cryomicroscopy. Nature 432:573 EP –. https://doi.org/10.1038/nature03079.

43. Dannhauser, P. N., and E. J. Ungewickell, 2012. Reconstitution of clathrin-coated bud and vesicle formation with minimal components. Nature Cell Biology 14:634–639. https://doi.org/10.1038/ncb2478.

44. Jin, A. J., K. Prasad, P. D. Smith, E. M. Lafer, and R. Nossal, 2006. Measuring the Elasticity of Clathrin-Coated Vesicles via Atomic Force Microscopy. Biophysical Journal 90:3333–3344. http://www.sciencedirect.com/science/article/pii/S0006349506725152.

45. Lherbette, M., L. RedlingshÃ¶fer, F. M. Brodsky, I. A. T. Schaap, and P. N. Dannhauser, 2019. The AP2 adaptor enhances clathrin coat stiffness. The FEBS Journal 286:4074–4085. https://febs.onlinelibrary.wiley.com/doi/abs/10.1111/febs.14961.

46. Sirotkin, V., J. Berro, K. Macmillan, L. Zhao, T. D. Pollard, and S. L. Schmid, 2010. Quantitative Analysis of the Mechanism of Endocytic Actin Patch Assembly and Disassembly in Fission Yeast. Molecular Biology of the Cell 21:2894–2904. https://doi.org/10.1091/mbc.e10-02-0157, pMID: 20587778.

47. Gallop, J. L., C. C. Jao, H. M. Kent, P. J. G. Butler, P. R. Evans, R. Langen, and H. T. McMahon, 2006. Mechanism of endophilin N-BAR domain-mediated membrane curvature. The EMBO Journal 25:2898–2910. https://www.embopress.org/doi/abs/10.1038/sj.emboj.7601174.

48. Henne, W. M., H. M. Kent, M. G. J. Ford, B. G. Hegde, O. Daumke, P. J. G. Butler, R. Mittal, R. Langen, P. R. Evans, and H. T. McMahon, 2007. Structure and Analysis of FCHo2 F-BAR Domain: A Dimerizing and Membrane Recruitment Module that Effects Membrane Curvature. Structure 15:839–852. https://doi.org/10.1016/j.str.2007.05.002.

49. Helfrich, W., 1973. Elastic properties of lipid bilayers: theory and possible experiments. Zeitschrift für Naturforschung C 28:693–703.

50. Derényi, I., F. Jülicher, and J. Prost, 2002. Formation and Interaction of Membrane Tubes. Phys. Rev. Lett. 88:238101. http://link.aps.org/doi/10.1103/PhysRevLett.88.238101.

51. Berro, J., V. Sirotkin, and T. D. Pollard, 2010. Mathematical Modeling of Endocytic Actin Patch Kinetics in Fission Yeast: Disassembly Requires Release of Actin Filament Fragments. Molecular Biology of the Cell 21:2905–2915. https://doi.org/10.1091/mbc.e10-06-0494, pMID: 20587776.

52. Footer, M. J., J. W. J. Kerssemakers, J. A. Theriot, and M. Dogterom, 2007. Direct measurement of force generation by actin filament polymerization using an optical trap. Proceedings of the National Academy of Sciences 104:2181–2186. https://www.pnas.org/content/104/7/2181.

53. Ma, R., and J. Berro, 2018. Structural organization and energy storage in crosslinked actin assemblies. PLOS Computational Biology 14:1–25. https://doi.org/10.1371/journal.pcbi.1006150.

54. Ma, R., and J. Berro, 2019. Crosslinking actin networks produces compressive force. Cytoskeleton 76:346–354. https://onlinelibrary.wiley.com/doi/abs/10.1002/cm.21552.

55. Jülicher, F., and U. Seifert, 1994. Shape equations for axisymmetric vesicles: A clarification. Phys. Rev. E 49:4728–4731. https://link.aps.org/doi/10.1103/PhysRevE.49.4728.

## REFERENCES

1. Derényi, I., F. Jülicher, and J. Prost, 2002. Formation and Interaction of Membrane Tubes. Phys. Rev. Lett. 88:238101. http://link.aps.org/doi/10.1103/PhysRevLett.88.238101.

2. Dmitrieff, S., and F. Nédélec, 2015. Membrane Mechanics of Endocytosis in Cells with Turgor. PLoS Comput Biol 11:1–15. http://dx.doi.org/10.1371%2Fjournal.pcbi.1004538.

